# Full title: Label free quantitative proteomics of *Qualea grandiflora* Mart. (Vochysiaceae) roots indicates an aluminium requirement for growth and development

**DOI:** 10.1101/396093

**Authors:** Natalia F. Cury, Renata C. C. Silva, Jessica R. Melo, Michelle S. F. Andre, Wagner Fontes, Carlos A. O. Ricart, Mariana S. Castro, Conceição E. S. Silveira, Marcelo V. de Sousa, Luiz A. R. Pereira

## Abstract

Aluminium in acid soils is a hindrance to crop growth. Nonetheless, Brazilian Cerrado possesses many species such as *Qualea grandiflora*, which are adapted to acid soils with large amounts of Al and accumulate this metal in its tissues and organs. Nonetheless, the mechanisms involved in these processes are poorly understood, mainly at molecular level. Thus, a root proteomic analysis was accomplished to identify Al-responsive proteins in *Q. grandiflora* plants. Concomitantly, a root growth analysis of plants grown with and without Al supplementation was conducted to determine the effects of Al on the whole plant. Subsequently, proteins from both treatments were identified and quantified by LC-MS/MS in a label-free fashion. From the 2,520 identified proteins, 410 were differentially abundant between the two treatments, which were associated with carbohydrate metabolism, redox activity, stress response and catabolism of organic compounds. Furthermore, Al was crucial for the growth and development of *Q. grandiflora*. In fact, this species may have an Al-dependent metabolism. Moreover, it was possible to correlate plant growth to Al-upregulated proteins that were directly involved in cell wall synthesis, oxidative phosphorylation, genetic information processing, and amino acid metabolism. Additionally, this work provides an extensive dataset of Al-regulated protein in *Q. grandiflora*, which will be crucial to understanding the functions of Al in this species.

## 1. Introduction

Aluminium (Al) is highly abundant in soils worldwide where it is found in different chemical species. In acidic conditions (pH < 5.5), these compounds are found as Al^+3^, which constitutes one of the most toxic form of Al, which is limiting for crop production. Nevertheless, there are plants that do not resent the presence of Al^+3^, and some of which are commonly found in the Brazilian Cerrado. The Cerrado biome is a neotropical savannah whose soils are usually acid, with low fertility and high Al contents [1,2]. Minerals play many crucial roles in plants. To determine the importance or essentiality of nutrients is necessary to verify the following criteria: 1) the element absence impairs the growth of the plant preventing it to complete its life cycle; 2) the signs of an element deficiency can only be reverted, exclusively, with its supplementation; 3) the element integrates a molecule or partakes of an essential biochemical reaction in the plant. These issues are termed as essentiality criteria. It has been suggested that some Cerrado plants, compulsorily, need Al to grow [3, 4]. Nonetheless, the importance of Al for these plants, including for *Qualea grandiflora* Mart., has not yet been determined.

The Cerrado encompasses about 2 million km^2^ and represents 23% of the Brazilian territory. This biome is regarded as the savannah with the highest biodiversity in the world and contains a large percentage of endemic plants [5]. Despite that, the Cerrado has been rapidly degraded, and its natural vegetation replaced by crops such as soybean, corn, and cotton. Therefore, the Cerrado is among the top 25 hotspots in the world [6]. Based on that, it is crucial to undergo efforts to study its native flora, which have enormous potential for medicinal, pharmaceutical, food as well as it constitutes a gene bank for plant genetic improvement.

As mentioned previously, several Cerrado plant species not only tolerate Al but also require it for proper growth and development. These plants usually belong to the families Rubiaceae, Melastomataceae, and Vochysiaceae. For instance, several species of *Miconia* spp (Melastomataceae), *Palicourea rigida* (Rubiaceae) and many plants of the genera *Vochysia* and *Qualea* (Vochysiaceae) need Al to grow and develop [3, 4]. Therefore, *Qualea grandiflora* Mart. (Vochysiaceae) has been chosen to investigate the metabolic role of this metal in native plants. Minerals play various crucial roles in plants. *Q. grandiflora* is a tree that reaches up to 15 m height and occurs in different phytophysiognomies of the Cerrado. This plant can accumulate about 5.0 g of Al.kg^-1^ of dry matter, and it is considered a compulsory Al-accumulating species [3].

Several genes and proteins associated with different metabolic pathways are induced in response to abiotic stimuli. The proteins encoded by the responsive genes to abiotic elicitors have also been located in various cell compartments such as cell wall, membranes, chloroplasts, and mitochondria. At the molecular level, there are many reports on Al-responsive genes in *Arabidopsis*, wheat, citrus, rice, soybean, tomato and many others plant species [7]. Besides, the information given by these studies is associated with Al tolerance/resistance in response to a stress elicited by this metal. For instance, in *Arabidopsis*, the absence of *STOP1* activity, a gene that encodes a zinc finger-like protein [8, 9], and *AtALMT1*, which encodes a malate transporter protein caused hypersensitivity to Al [10].

Also, in *Arabidopsis*, increased expression of *WAK1*, a gene encoding a cell wall kinase, significantly enhances Al tolerance [11]. Moreover, it has been reported that protein interacts with AtGRP3, a glycine-rich protein to control root size [12]. In fact, the outcome of AtWAK1/AtGRP3 interaction is usually cell elongation repression, which consequently results in shorter roots [12]. It has been proposed that only free AtWAK1 (no AtGRP3 interaction) augments Al tolerance in *Arabidopsis* [11, 12]. As consequence, *Arabidopsis grp3-1* knockout mutants have longer roots and a higher tolerance to Al, mimicking the overexpression of *AtWAK1* gene [11, 12].

Furthermore, proteomics has been used to identify Al-responsive proteins. For example, Wang et al. (2014) found 106 differentially abundant proteins from Al-treated rice roots and observed that the glycolysis/gluconeogenesis processes were upregulated by Al. In addition, all glycolysis/gluconeogenesis related genes were more expressed in Al-tolerant rice cultivar than in the sensitive [13]. This fact suggests energy availability is crucial for Al-tolerance.

To date and to the best of our knowledge, data on genes and proteins involved in Al metabolism and accumulation from Cerrado plants are rare, and there is no proteomic study in plants where the lack of Al maybe stressful. Therefore, the present investigation is part of an ongoing research project directed towards unravelling the role of Al in native species. Accordingly, *Q. grandiflora* plants were grown with and without Al supplementation with the purpose of quantifying the abundance of Al-responsive proteins. The differential abundance of proteins allowed the drawing of an initial picture of which metabolic pathways are either up or downregulated due to the presence of this metal.

## 2. Material and Methods

### 2.1. Experimental design

A flowchart was elaborated indicating the main steps carried out in the growth and proteomic analyses of roots of *Q. grandiflora* in response to Al (Fig 1).

**Fig 1.**
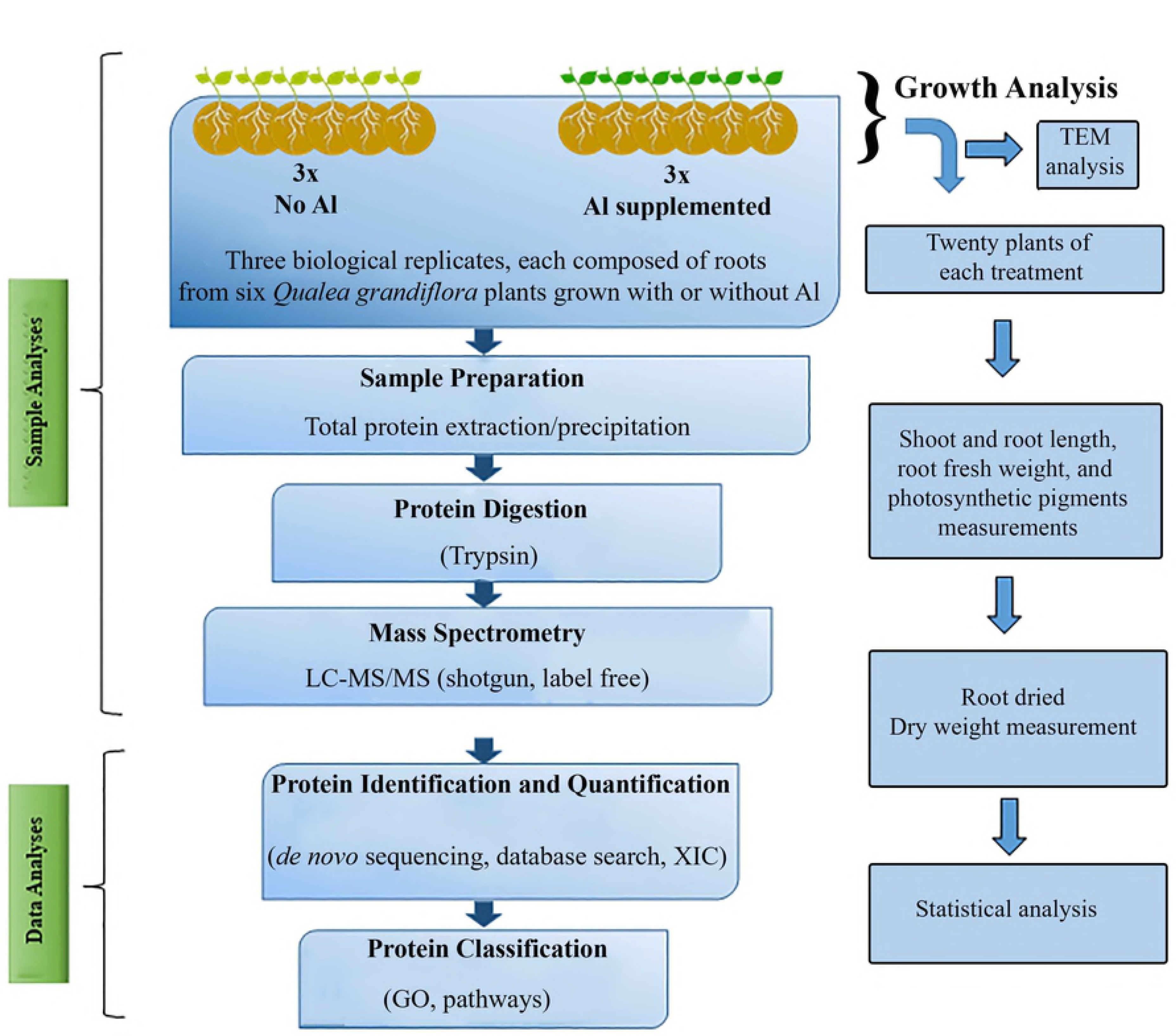
A chart of *Q. grandiflora* root proteomic, growth, and transmission electron microscopy analyses. Both analyses were accomplished using four-month-old *Q. grandiflora* plants grown with or without Al supplementation.

### 2.2. Plant materials, growth conditions and treatments

*Qualea grandiflora* Mart. (Vochysiaceae) seeds were sterilized, put to germinate in a germination chamber and then cultivate in sterile vermiculite. Forty *Q. grandiflora* plants (20 per treatment) were watered every two days with 1/5 MS (Murashige and Skoog) salts [14], pH 4.2-4.5, supplemented or not with AlCl_3_ (150 µM). Root samples of *Q. grandiflora* from Al-treated and no-Al treatments were collected after 90 days of cultivation. Each biological replicate was composed of the whole root system from six *Q. grandiflora* plants, and three biological replicates of each treatment were analysed. Subsequently, the samples were immediately frozen in liquid nitrogen and kept in a freezer at −80 °C for protein extraction.

### 2.3. Growth analysis

For growth analysis, each treatment was composed of 20 *Q. grandiflora* plants grown with or without Al in three replicates. At 120^th^ day of cultivation, the following parameters were analysed: shoot length, root length, root fresh and dry weights, and photosynthetic pigment contents. For fresh weight, *Q. grandiflora* plants were removed from vermiculite, slightly dried on paper towel and weighed. For dry weight, roots were dried in an oven at 72 °C, for 72 h and then weighed.

Six leaf discs of 0.5 cm in diameter from three different plants of each treatment were placed in microcentrifuge Eppendorf tubes containing 2 ml DMF (N, N -dimethylformamide-HCON (CH_3_) _2_). The tubes were wrapped in aluminium foil and stored for 48 h at 4 °C. Afterwards, the absorbance of each extract was measured at the wavelengths of 663.8 nm, 646.8 nm and 480 nm on a Thermo Spectronic spectrophotometer. Therefore, the contents of chlorophylls a and b, and carotenoids was calculated as proposed by Wellburn (1994), as follows:

1-Chlorophyll a = ((12*Ab663.8-3.11*Ab646.8)*Vol)/A)
2-Chlorophyll b = ((20.78*Ab646.8 - 4.88*Ab663.8)*Vol)/A)
3-Carotenoids = (((1000*Chlorophyll a-1.12*Chlorophyll b-34.07*Ab480)/245)*2)/A)

#### 2.3.1. Statistical Analysis

All growth analysis data was statistically analysed by two-way ANOVA and the differences among the means were tested by the Student’s t-test (p< 0.05).

### 2.4 Transmission Electron Microscopy

*Q. grandiflora* leaf samples of plants cultivated for 120 days from both treatments were fixed overnight in Karnovsky fixative in 0.05 M sodium cacodylate buffer (pH 7.3) at 4 °C. The leaf samples were: green leaves from Al-treated plants, green leaves and yellow leaves from non-treated plants. Subsequently, the leaf samples were post-fixed in 1% osmium tetroxide (OsO_4_) in the dark for 30 min. Then, the samples were dehydrated in a graded acetone series (30%, 50%, 70%, 90%, 100%). Afterwards, the samples were infiltrated in acetone:SPURR (v/v) and embedded in SPURR (100%) resin for 24 h each step. Ultrathin sections (60 nm) were made on an ultramicrotome and then mounted on nickel grids (300 mesh) and observed on the transmission electron microscope (Jeol JEM 1011) at 60 KV.

### 2.5. Root protein extraction

Approximately 100 mg of each root sample were ground in liquid nitrogen using a mortar and pestle. Then, the powder was added to a solution of 10% (w/v) trichloroacetic acid and 0.07% (v/v) β-mercaptoethanol in cold acetone, and the resulting suspension was thoroughly mixed by vortexing, and then incubated for 3 h at 4 °C.

After incubation, samples were centrifuged at 10,000 g for 20 min at 4 °C. Then, the supernatant was removed, and the remaining pellet was washed five times with 10% (w/v) trichloroacetic acid in acetone until the total disappearance of pigments. The pellet was dried using a Speed-Vac concentrator and resuspended in rehydration buffer (7 M urea, 2 M thiourea, 250 mM TEAB, pH 8.5). Protein concentration was determined by using Qubit^®^ 2.0 assay (Invitrogen, Carlsbad, USA) and extracted protein quality was assessed in 10% SDS-PAGE.

### 2.6. Protein digestion

The extracted proteins (200 μg) were reduced with 10 mM dithiothreitol (DTT) for 60 min at 56 °C and alkylated with 100 mM iodoacetamide (IAA) for 60 min at 37 °C in the dark. Then, the samples were diluted in 100 mM NH_4_HCO_3_ (ammonium bicarbonate), pH 8.1. Subsequently, the alkylated proteins were digested with trypsin (1:50 v/v – Promega, Madison, USA) at 37 °C for 16 h. After digestion, the resulting peptide solution was acidified with 0.1% trifluoroacetic acid and centrifuged at 10,000 g for 10 min. Then, the supernatant was desalted in homemade microcolumns (C_18_ stage tips), vacuum-dried, and resuspended in 0.1% (v/v) formic acid. After, the sample was quantified by Qubit^®^ 2.0 (Invitrogen, Carlsbad, USA) for further analysis by liquid nano-chromatography coupled to a mass spectrometer.

### 2.7. Chromatography and nano LC-MS/MS analysis

The tryptic peptides were applied to an Ultimate 3000 liquid chromatographer (Sunnyvale, USA) for reversed phase nano-chromatography as following: three technical replicates of 1 μg from each biological replicate were injected into a trap column (2 cm × 100 μm), containing C_18_ 5 μm particles. The samples were eluted to an analytical column (32 cm × 75 μm, C_18_ 3 μm), and then to the ionization source of the spectrometer. The elution gradient was composed of 0.1% (v/v) formic acid in water (solvent A), and 0.1% (v/v) formic acid in acetonitrile (solvent B), in a gradient of 2% to 35% solvent B for 180 min.

The eluted fractions were placed in the ionization source of the LTQ Orbitrap Elite mass spectrometer (Thermo Fisher Scientific, Germany) and were analysed in DDA (data dependent acquisition) mode, produced MS1 spectra in the Orbitrap analyser (resolution of 120,000 FWHM at 400 m/z) in the range of 300-1650 m/z. For each MS1 spectrum, the 15 most intense ions with charge greater than 2 were automatically chosen and directed to high-energy collision-induced dissociation (HCD) fragmentation. Repeated fragmentation of the same precursor was prevented by dynamic exclusion for 90 s. The configuration for HCD was: insulation window 2.0 m/z, automatic gain control (AGC) of 5 × 10^6^ and maximum fill time (maximum TI) of 100 ms, with collision energy normalized to 35% and threshold for the selection of 3000.

### 2.8. Database search and label-free quantification

The files obtained from the mass spectrometer were analysed using the software Progenesis IQ for alignment of the MS1 peaks found in the chromatograms, extracted ion chromatogram (XIC)-based quantification and normalization. A first statistical analysis was performed before the identification of the MS1 features, to filter for identification only those presenting ANOVA p-values < 0.05.

After the peptide peaks were quantified and grouped, the identification of proteins was performed using Peaks 7.0 software, which deduced sequences from the fragmentation information and searched the sequence database of the *Q. grandiflora* transcriptome deposited at the NCBI (National Centre for Biotechnology Information), BioProject PRJNA358394 (130.704 *Searched Entries*). The search parameters used were: precursor ion mass error tolerance of 10 ppm, MS/MS mass tolerance of 0.05 Da, carbamidomethylation of cysteine residues (fixed modification) and deamidation and methionine oxidation (variable modification). Trypsin was selected as the digestion enzyme, and up to two missed cleavage sites per peptide were allowed.

The identified proteins were filtered at a rate of 1% for false discovery rate (FDR), and a minimum of one unique peptide per protein was required for identification.

The protein identification information was imported into the Progenesis IQ software, which combined them with previously generated quantitative data. Multivariate PCA analysis was performed in Progenesis to evaluate the grouping of replicates and conditions, as well as to cluster the abundance profiles. In this study, a protein was considered differentially abundant when it presented fold-change ≥ 1.5 with p ≤ 0.05 after the ANOVA test at the protein level.

### 2.9. Protein classification and identification of Al-responsive pathways

For functional classification, the differentially abundant proteins in *Q*. *grandiflora* were analysed using the Blast2GO 4.1 Gene Ontology (GO) analysis tool and assigned to the three GO concepts: biological processes, molecular function and cellular components.

Metabolic pathways analysis was performed by searching the KEGG (Kyoto Encyclopedia of Genes and Genomes) database using the BlastKOALA (KEGG Orthology And Links Annotation) server [**15**] for functional annotation. The following parameters were applied: plant taxonomic group and eukaryotes family database as reference data, as well as to verify the overlapping enrichment between data of the present study and the pathways present in the database.

### 2.10. Analysis of protein interactions

The proteins identified as differentially abundant were also investigated for their interaction with other proteins using bioinformatics tools such as STRING 10.0 (Search Tool for the Retrieval of Interacting Genes / Proteins) program accessible at http://string-Db.org [16]. The STRING 10.0 analysis was performed using the highest confidence score (0.900) to compare with homologous proteins of *Arabidopsis thaliana* database.

## 3. RESULTS

### 3.1. The lack of Al resulted in depletion of *Q. grandiflora* growth and development

#### 3.1.1. Photosynthetic pigment contents Photosynthetic apparatus

The quantification of the photosynthetic pigments showed significant differences between Al-supplemented and non-supplemented *Q. grandiflora* plants. Plants grown with Al had 45-50% higher contents of chlorophyll *a, b*, and carotenes (Table 1). This data is consistent with the overall appearance of the plants. Morphologically, the lack of Al in the nutritional solution resulted in chlorotic plants, while the plants treated with Al the leaves were green (Fig 2). These facts did indicate that plants grown without Al may have resented its absence.

**Table 1.** Effect of Al supplementation on photosynthetic pigments (chlorophyll *a*, chlorophyll *b*, and carotenes) content (mg.cm^-2^) of 120-day-old *Q. grandiflora* plants.

| Treatment | Chlorophyll *a* (± SD) | Chlorophyll *b* (± SD) | Total Carotenoids (± SD) |
| --- | --- | --- | --- |
| With Al | 24.73 ± (2.91) **b** | 8.27 ± (0.32) **b** | 6.20 ± (0.22) **b** |
| No Al | 11.95 ± (4.34) **a** | 4.64 ± (1.00) **a** | 3.52 ± (0.77) **a** |

The means followed by the same letters within same column do not differ statistically by the student's *t* test (*p* < 0.05), *n* = 5. SD: standard deviation.

**Fig 2.**
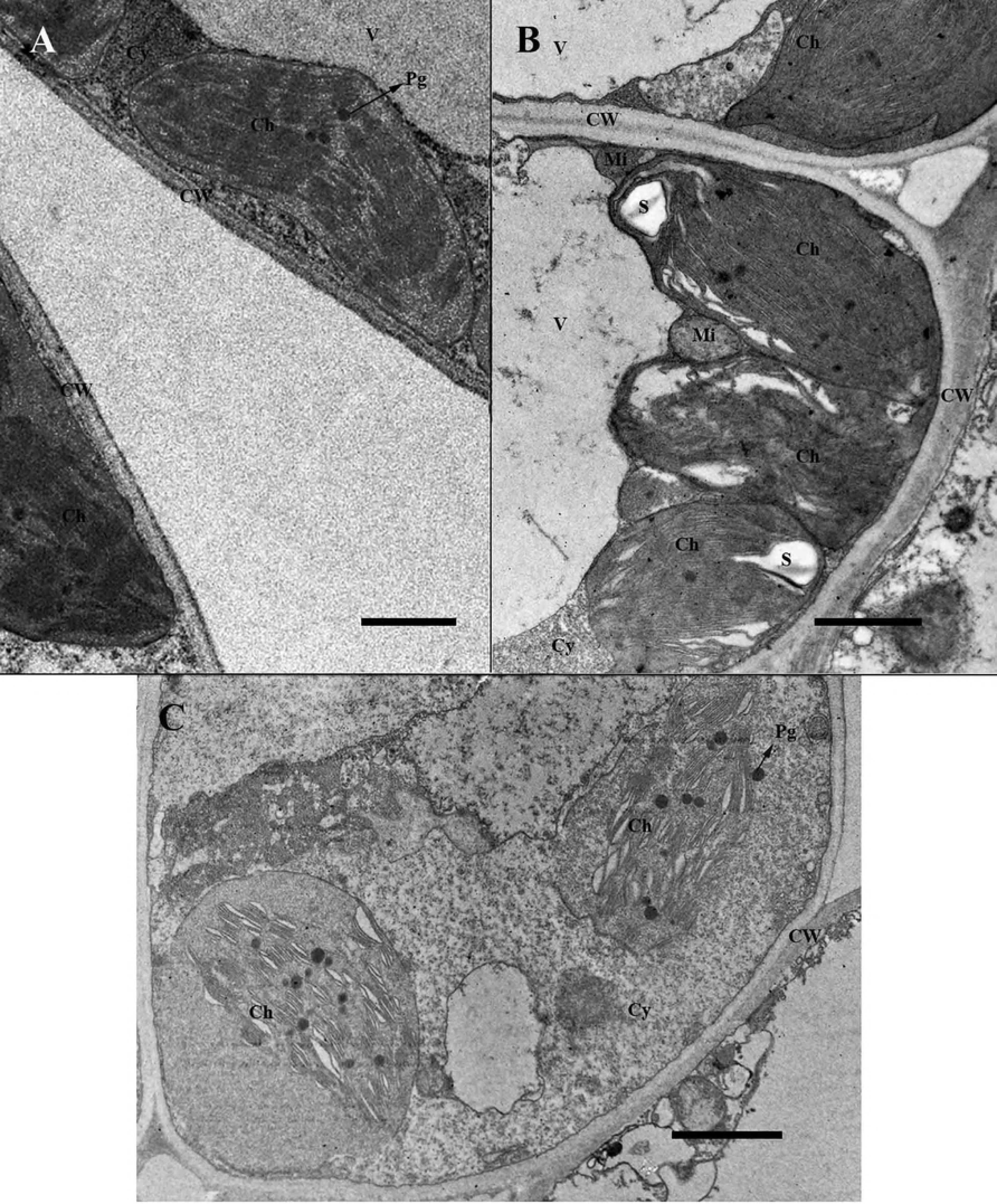
Morphological aspects of *Q. grandiflora* plants grown with or without Al supplementation. A) Upper view of an Al-treated plant showing the healthy aspect of leaves. B) Roots from plants that received Al supplementation. C) Upper view of a *Q. grandiflora* plant not treated with Al showing signs of chlorosis. D) Roots from non-treated plants. Scale bar: A-C = 1.5 cm; B-D = 1.7 cm.

Furthermore, Figure 3 depicts ultrastructural features of chloroplasts from leaves of *Q. grandiflora* plants grown with or without Al. It is noteworthy that the chloroplasts from each treatment had a distinct structure (Fig 3). Note that the chloroplast from leaves of Al-supplemented plants had a standard structure, with typical shape and regular internal membrane system with grana and thylakoids (Fig 3 A). Differently, the chloroplasts from non-treated plants showed abnormal structure. Moreover, even the chloroplasts from green leaves already had a peculiar internal membrane system (Fig 3 B). Note that in these chloroplasts the lumen of thylakoids appeared dilated, and no typical assembly of granum (Fig 3 B). Furthermore, in yellowish leaves, the thylakoid dilation was very pronounced, and the stroma appeared to be exceedingly large compared with chloroplasts from green leaves (Fig 3 B).

**Fig 3.**
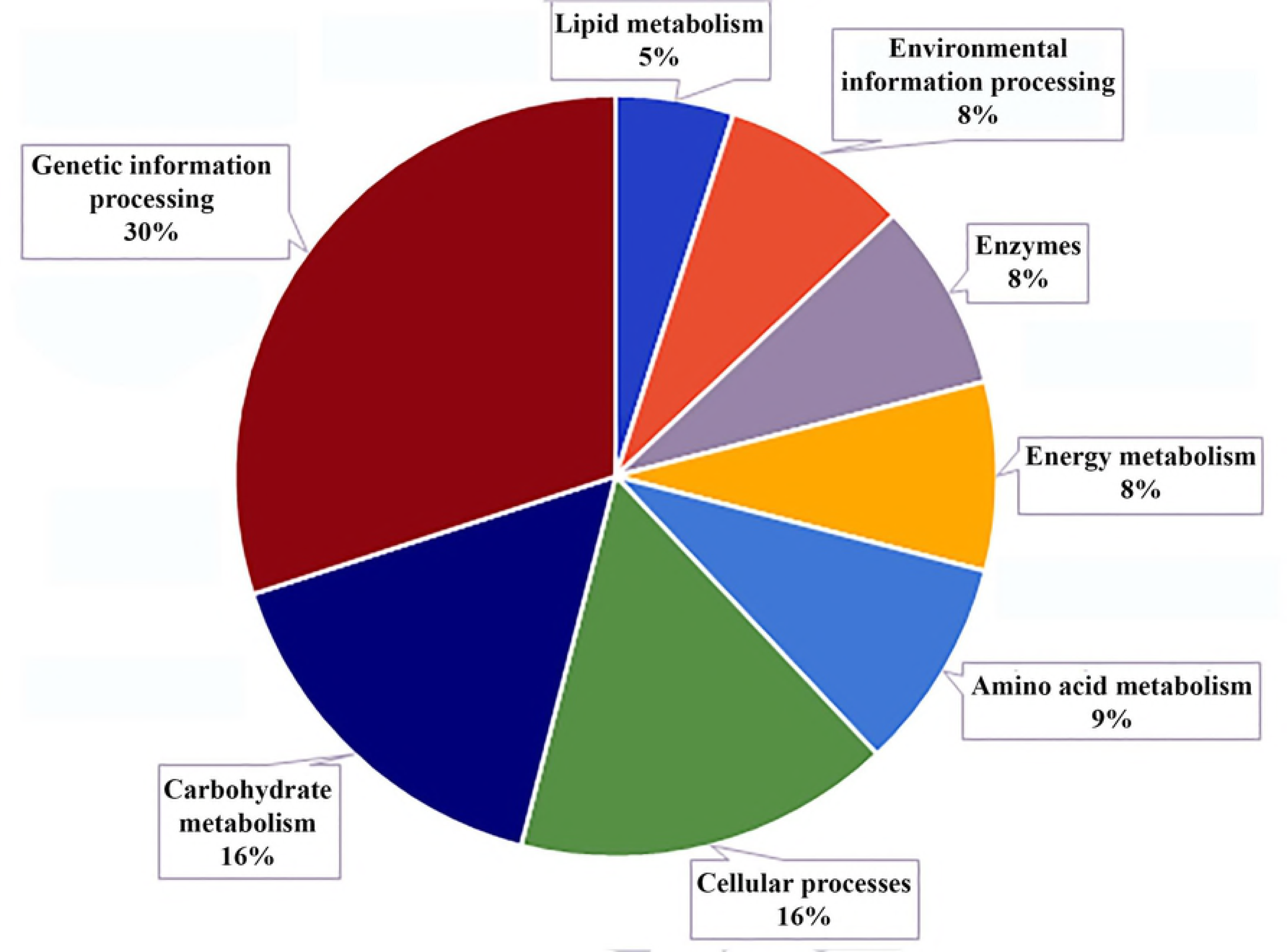
Transmission electron micrographs of *Q. grandiflora* chloroplasts from plants grown with or without Al supplementation. A) Chloroplasts from green leaves of an Al-treated plant. B) Chloroplasts from green leaves of a plant that was not supplemented with Al. C) Chloroplasts from a yellowish *Q. grandiflora* leaf of a plant not treated with Al. Ch: chloroplast. CW: cell wall. Cy: cytoplasm. Mi. mitochondrion. Pg: plastoglobule. S: starch. V: vacuole. Scale bar: 0.5 µm

#### 3.1.2. Al Influence on Shoot and Root Growth

A growth analysis of *Q. grandiflora* shoots and roots was performed to investigate whether the morphological differences observed between the treatments were associated with the lack of Al supplementation. The results indicated that Al was critical for growth and development of *Q. grandiflora* plants. Note that leaves from Al-supplemented plants were greener than those from non-treated plants. Moreover, it is noteworthy that shoots and roots from Al-treated plants were considerably higher and the roots were longer and more branched (Fig 2, Table 2). Furthermore, after 120 days of cultivation the average length of shoot of Al-treated plants was about 14% higher than shoots from non-treated plants. In roots this difference was even greater and reached 24% in favour of plants grown with Al (Table 2).

**Table 2.** Root average length (mm) and growth rates (%) of *Qualea grandiflora* plants grown in the presence and absence of Al.

| Treatment | *Qualea grandiflora* shoot length (mm) | *Qualea grandiflora* root length (mm) |
| --- | --- | --- |
| $\frac{1}{5}$ MS + AlCl<sub>3</sub> (150 μM) | 79.61 (±2.09) A | 159.76 (±5.48) A |
| $\frac{1}{5}$ MS | 68.18 (±2.78) B | 121.48 (±3.90) B |

Numbers followed by the same letters, within the same column and, are statistically similar as calculated by the Student's t-test ( $p < 0.05$ ), $n=20$ . Numeric values the means ± standard error.

Besides, root biomass accumulation was determined by measuring root fresh and dry weights (Table 3).

**Table 3.**
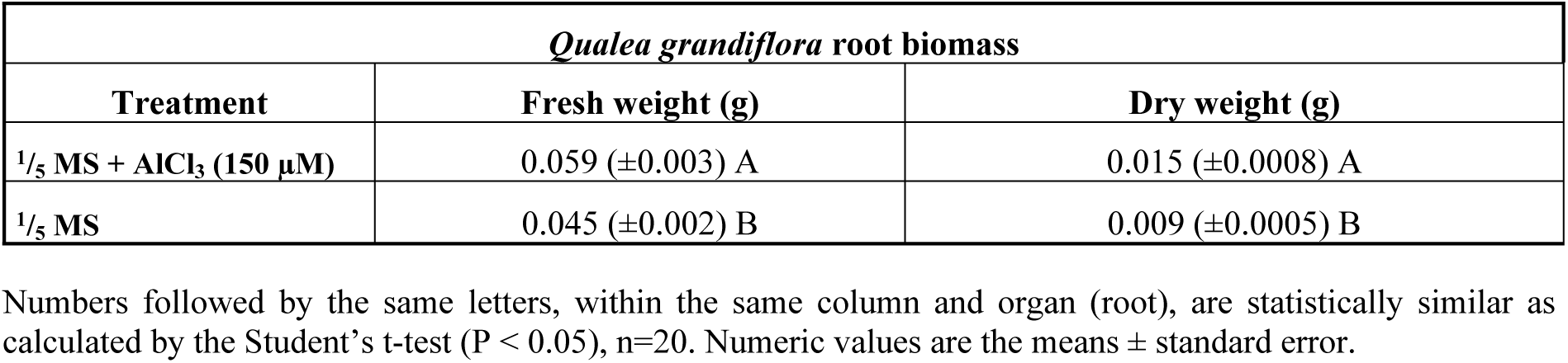
Average of root biomass (g) of *Qualea grandiflora* plants grown with or without Al after 120 days of cultivation.

#### 3.1.3 Root biomass accumulation

Consistent with root length data, Al-treated plants had significantly higher root biomass compared with those not treated (Table 3). The results showed that after 120 days of cultivation root fresh and dry weights of roots grown with Al were respectively 24% and 40% higher than those that did not receive Al. This fact indicates that the root system of *Q. grandiflora* was stimulated by Al.

### 3.2. Identification of Al-responsive proteins in *Q. grandiflora* roots

Gel-free and label-free approach were used to determine the proteomic changes in *Q. grandiflora* roots grown with or without Al-supplementation. A total of 2,520 distinct proteins were identified in both treatments by using a transcriptome sequence database and PEAKS and PepExplorer software. For the identification of differentially abundant proteins, it has been considered a fold-change ≥ 1.5 and a p-value ≤ 0.05 (Supplementary Table 1). Thus, in *Q. grandiflora* roots a total of 410 differentially abundant Al-responsive proteins were identified (Supplementary Table 2), and most of which upregulated, totalizing 274 proteins (67%). Complementarily, the remaining 136 (33%) were downregulated in response to Al.

The differentially abundant proteins were catalogued according to their functional categories established by GO terms (Blast2GO), and the proteins are depicted with individual identification, the number of peptides, score and fold change values (Supplementary Table 2).

### 3.3. Functional classification of Al-responsive proteins

The 410 Al-regulated proteins were analysed and classified within the following GO terms as determined by Blast2GO 4.1: biological processes (BP), cellular components (CC) and molecular functions (MF). It is important to say that the actual number of Al-responsive proteins may be different from the summation of all categories, since some proteins can be classified into multiples groups.

Figure 4 depicts the main biological processes induced by Al in roots of *Q. grandiflora*. Therefore, some proteins involved in response to stimulus (7%), biological regulation (4%), metabolic process (28%), cellular process (34%), single cellular process (23%), and cellular component organization/biogenesis (4%) were differentially abundant.

**Fig 4.**
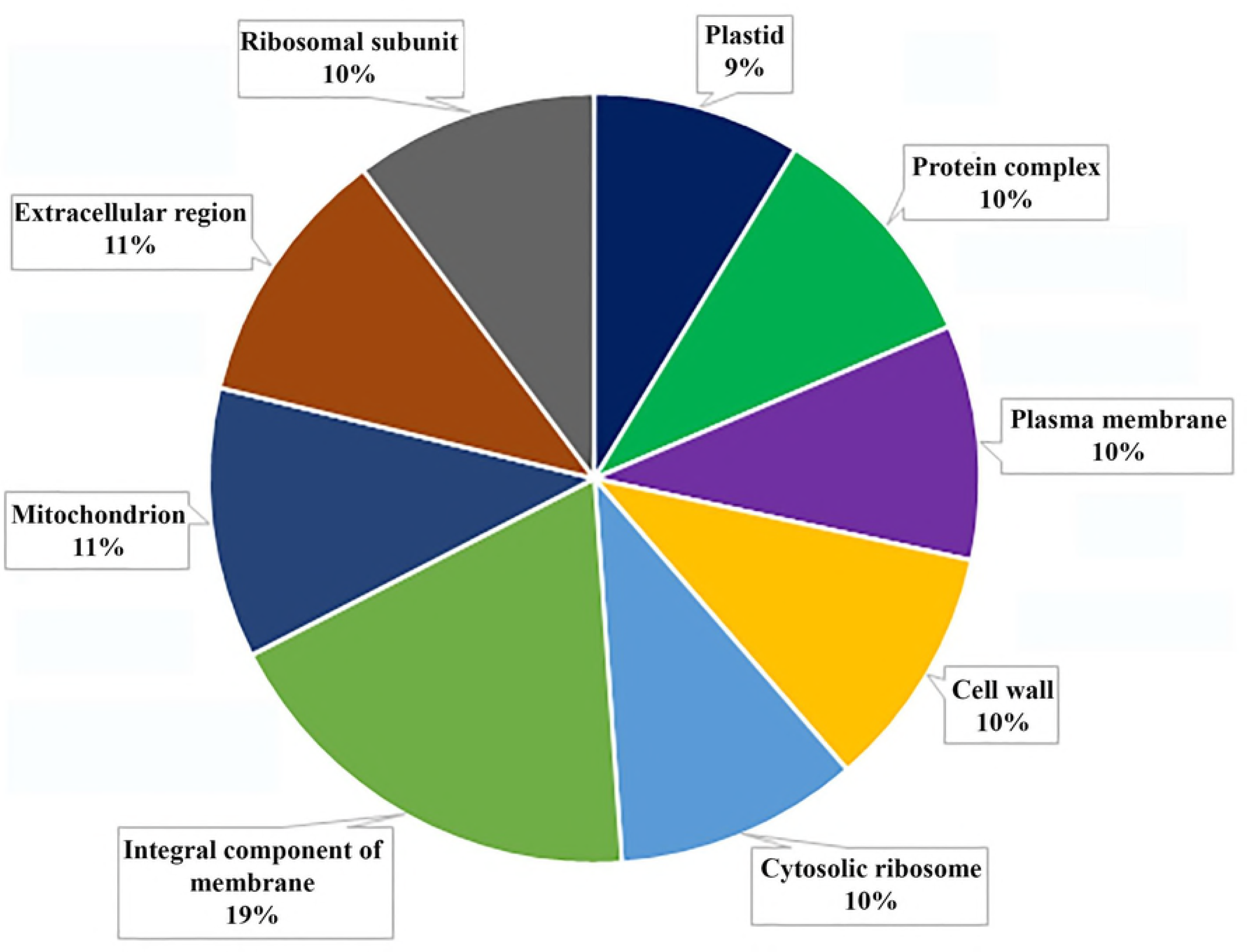
Biological processes responsive to Al in *Qualea grandiflora* plants. Distribution of differentially abundant proteins responsive to Al and respective biological processes. The numbers represent the occurrence in each GO term.

The most enriched BP categories in *Q. grandiflora* roots were metabolic processes whose proteins were associated with nitrogen compound metabolism, biosynthetic process, organic substance metabolism, catabolic process, regulation of cellular process, single organism metabolism, and primary metabolism. In fact, within these categories the primary metabolism had 112 upregulated proteins in roots from Al-treated plants (Supplementary Table 2).

In addition, the main cellular components associated with Al response in *Q. grandiflora* roots were located at plasma membrane, cell wall, plastid, mitochondrion, and ribosomal subunits, as well as the extracellular compartment (Fig 5).

**Fig 5.**
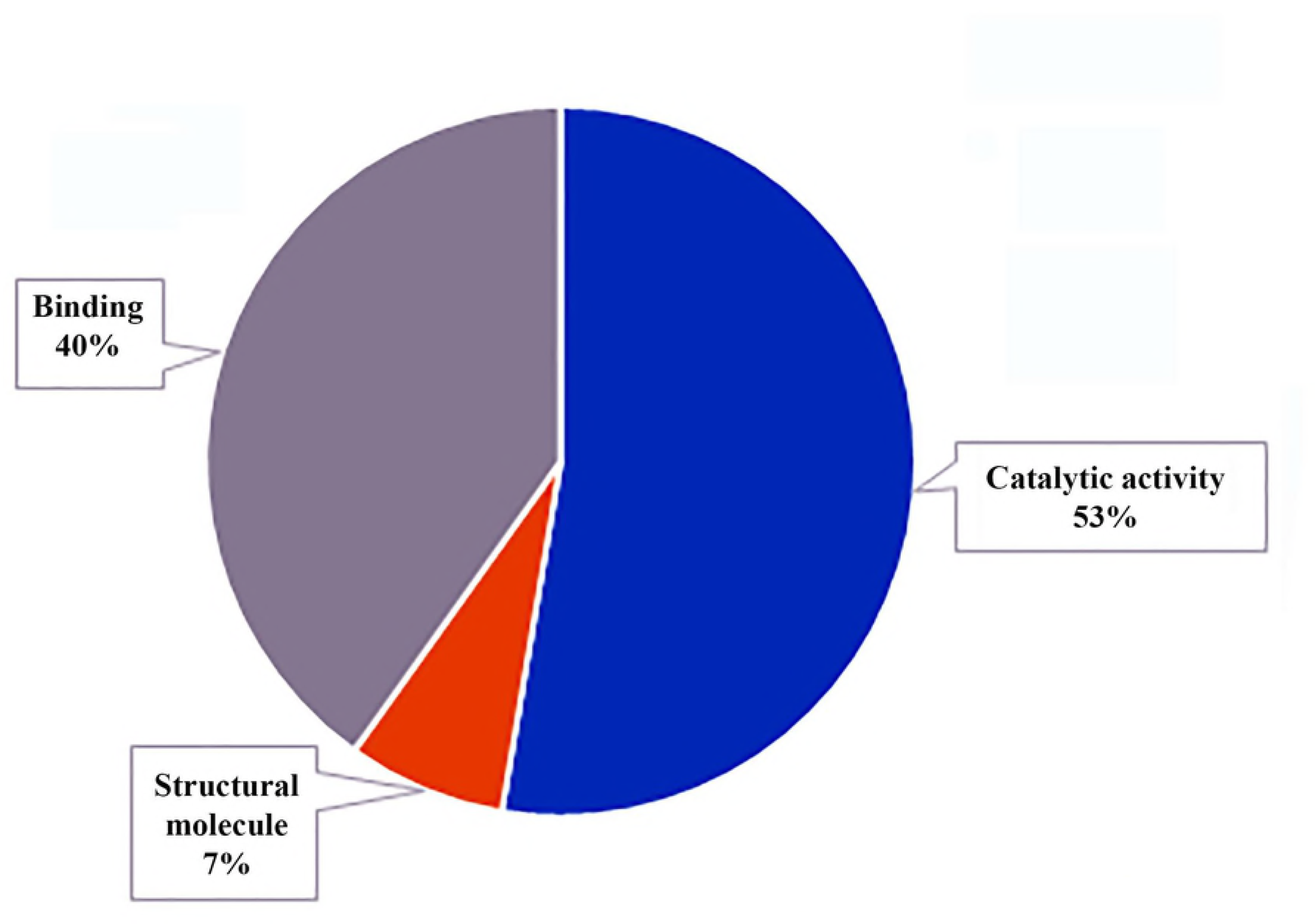
Chart showing the distribution of differentially abundant proteins responsive to Al and their association with cellular components in *Qualea grandiflora* plants according to GO term analysis.

Concerning the molecular function category of Al-responsive proteins, most of them were associated with binding activity (40%), catalytic activity (53%), and structural molecule activity (7%) as shown in Figure 6.

**Fig 6.**
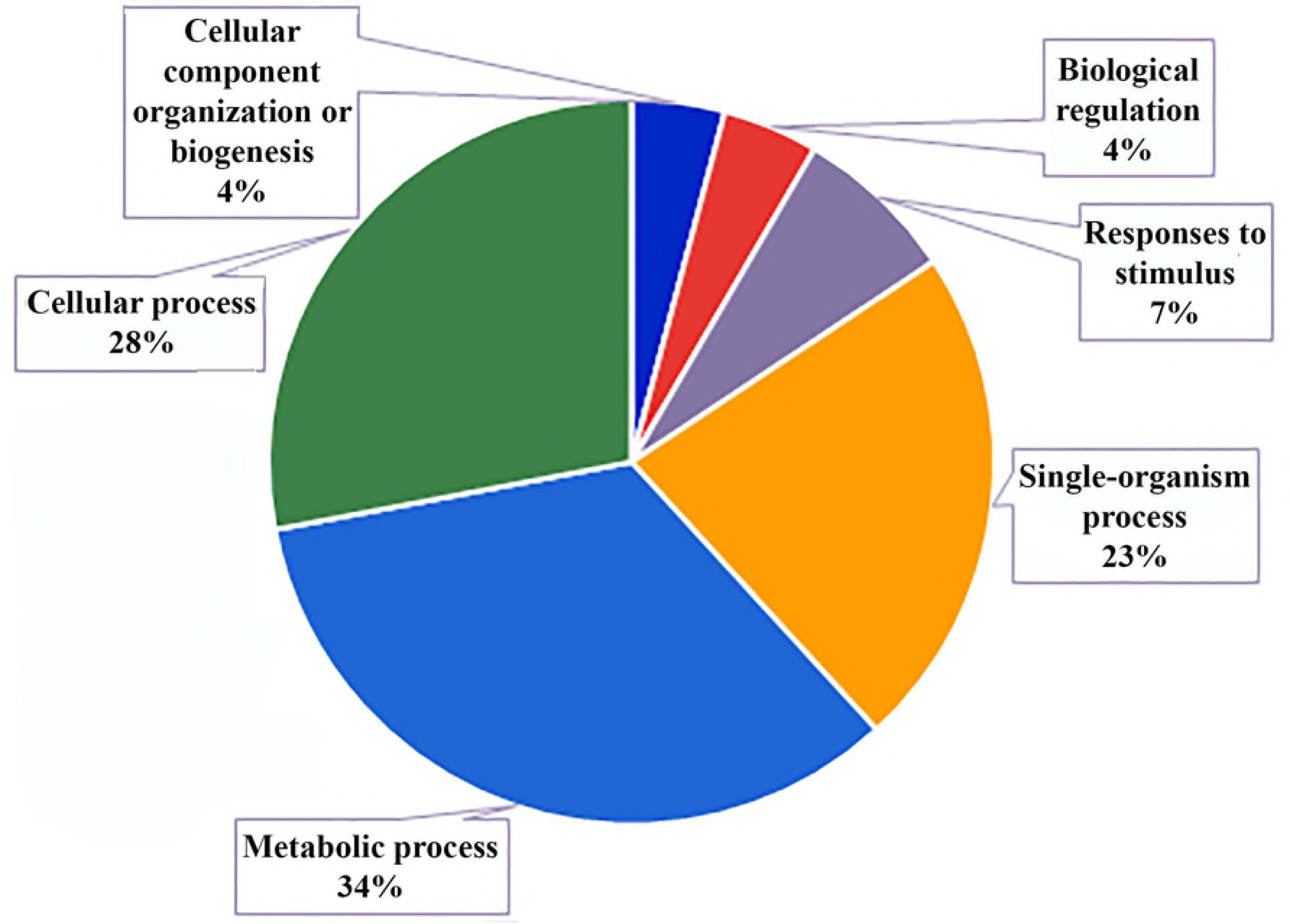
Main molecular functions of proteins involved in the response to Al in *Qualea grandiflora* plants.

Within binding activity, protein binding, metal ion binding, ATP binding and nucleic acid binding were the most common. Furthermore, proteins with hydrolase, redox, and transferase activities were also found to be differentially abundant in roots of *Q. grandiflora*. The smaller group in this category was the structural function.

### 3.4. Metabolic pathways upregulated by Al in roots of *Qualea grandiflora*

To determine which metabolic pathways were most responsive to Al, differentially abundant proteins were identified in roots of *Q. grandiflora* using the KEGG (Kyoto Encyclopedia of Genes and Genomes) database. Compared with no Al treatment, 272 (66.3%) out the 410 differentially abundant proteins were annotated in the KEGG database and assigned to 117 different metabolic pathways. Also, the results showed that differentially abundant proteins were mainly involved in genetic information processing, followed by carbohydrate metabolism, cellular processes, amino acid metabolism, energy metabolism and lipids (Fig 7).

**Fig 7.**
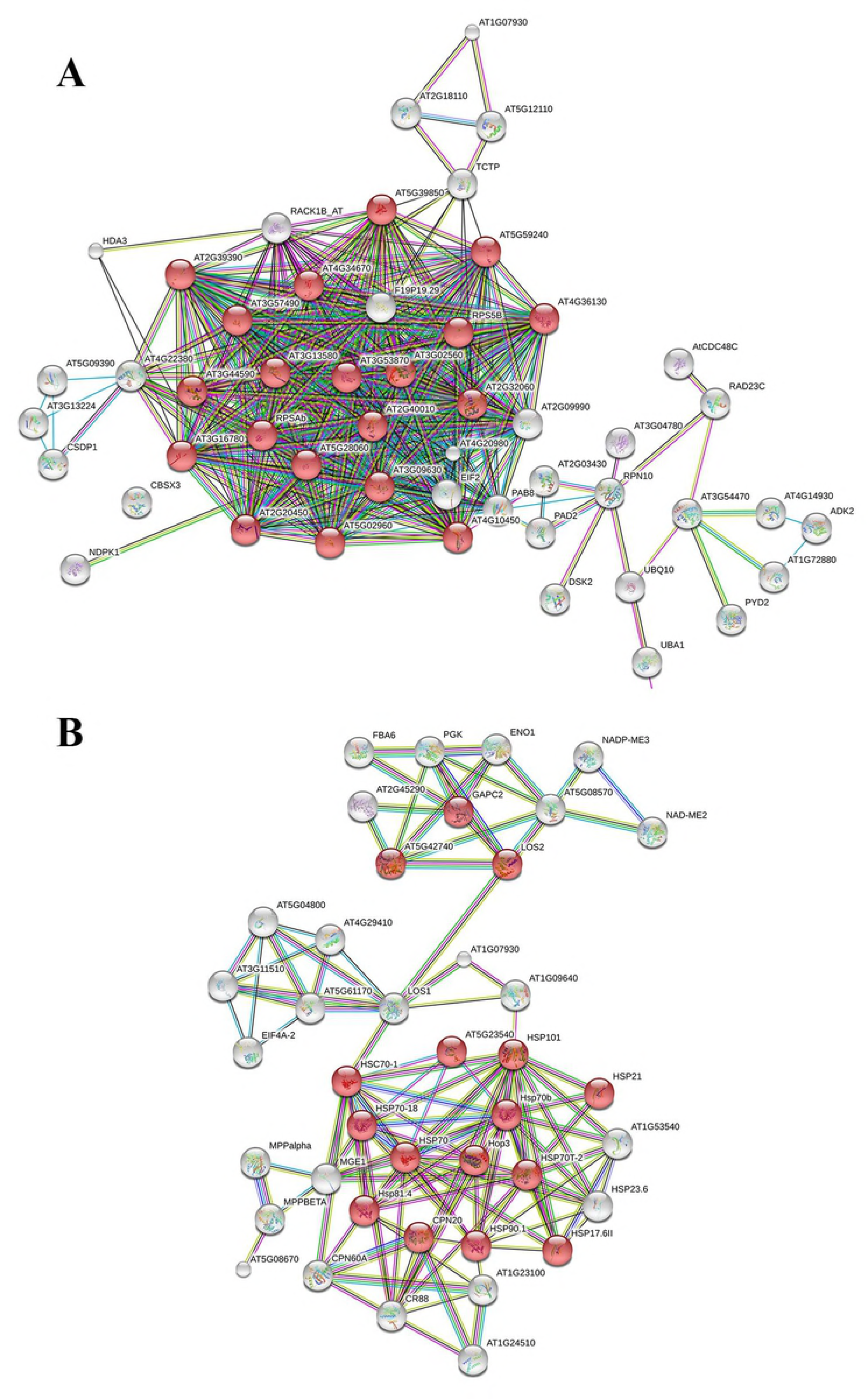
Main metabolic pathway categories in which the differentially abundant Al responsive *Qualea grandiflora* root proteins were associated with during KEGG analysis.

As the number of metabolic pathways involved in Al metabolism of *Q. grandiflora* roots was considerably high (117), an enrichment analysis (STRING) was made to determine which metabolic routes were the most relevant (Table 4).

**Table 4.** Enrichment analysis (STRING) of metabolic pathways associated with *Qualea grandiflora* root differentially abundant Al responsive proteins using KEGG database.

|  | Pathway ID | Pathway description | Number of proteins | FDR |
| --- | --- | --- | --- | --- |
| 1 | 3010 | Ribosome | 21 | 1.36e-11 |
| 2 | 20 | Citrate cycle (TCA cycle) | 8 | 9.4e-07 |
| 3 | 630 | Glyoxylate metabolism | 8 | 9.4e-07 |
| 4 | 1230 | Biosynthesis of amino acids | 14 | 9.4e-07 |
| 5 | 71 | Fatty acid degradation | 7 | 1.11e-06 |
| 6 | 1212 | Fatty acid metabolism | 7 | 6.74e-05 |
| 7 | 240 | Pyrimidine metabolism | 7 | 0.000814 |
| 8 | 592 | alpha-Linolenic acid metabolism | 4 | 0.00141 |
| 9 | 270 | Cysteine and methionine metabolism | 6 | 0.00209 |
| 10 | 51 | Fructose and mannose metabolism | 4 | 0.0057 |
| 11 | 640 | Propanoate metabolism | 3 | 0.0105 |
| 12 | 603 | Glycosphingolipid biosynthesis - globo series | 2 | 0.0146 |
| 13 | 260 | Glycine, serine and threonine metabolism | 4 | 0.0164 |
| 14 | 1220 | Degradation of aromatic compounds | 2 | 0.0164 |
| 15 | 760 | Nicotinate and nicotinamide metabolism | 2 | 0.0261 |
| 16 | 4145 | Phagosome | 4 | 0.0261 |
| 17 | 780 | Biotin metabolism | 2 | 0.0291 |
| 18 | 280 | Valine, leucine and isoleucine degradation | 3 | 0.0327 |
| 19 | 480 | Glutathione metabolism | 4 | 0.0419 |
| 20 | 190 | Oxidative phosphorylation | 5 | 0.0425 |
| 21 | 450 | Seleno compound metabolism | 2 | 0.0425 |
| 22 | 30 | Pentose phosphate pathway | 3 | 0.0434 |
| 23 | 230 | Purine metabolism | 5 | 0.0457 |
| 24 | 670 | One carbon pool by folate | 2 | 0.0481 |

Therefore, the enrichment analysis revealed 24 metabolic routes that were statistically significant (FDR<0.05) in *Q. grandiflora* roots in response to Al (Table 4). Moreover, the most relevant metabolic pathways indicated that proteins with differential abundance were mainly involved in the metabolism of glyoxylate (N = 8), oxidative phosphorylation (N = 5), fatty acid degradation (N = 7), metabolism of purine and pyrimidine (N = 12), metabolism of methionine and cysteine (N = 6), and in ribosomal activity (N = 21).

### 3.5. Protein-protein interactions

Protein-protein interactions may play a crucial role in cellular function. Hence, to understand the complex relationships of protein interactions in roots of *Q. grandiflora*, a STRING analysis was performed with a high level of stringency (confidence score = 0.900).

A network of interaction between ribosomal family proteins involving 21 proteins whose abundance was significantly increased with FDR 1.36 e-11 (Fig 8A) was observed. The STRING analysis also indicated that the differential proteins with significantly decreased abundance and a strong interaction in response to stresses were present. This interaction network involved 31 proteins, mostly HSPs (heat shock proteins) with FDR 1.07 e-05 (Fig 8B).

**Fig 8.**
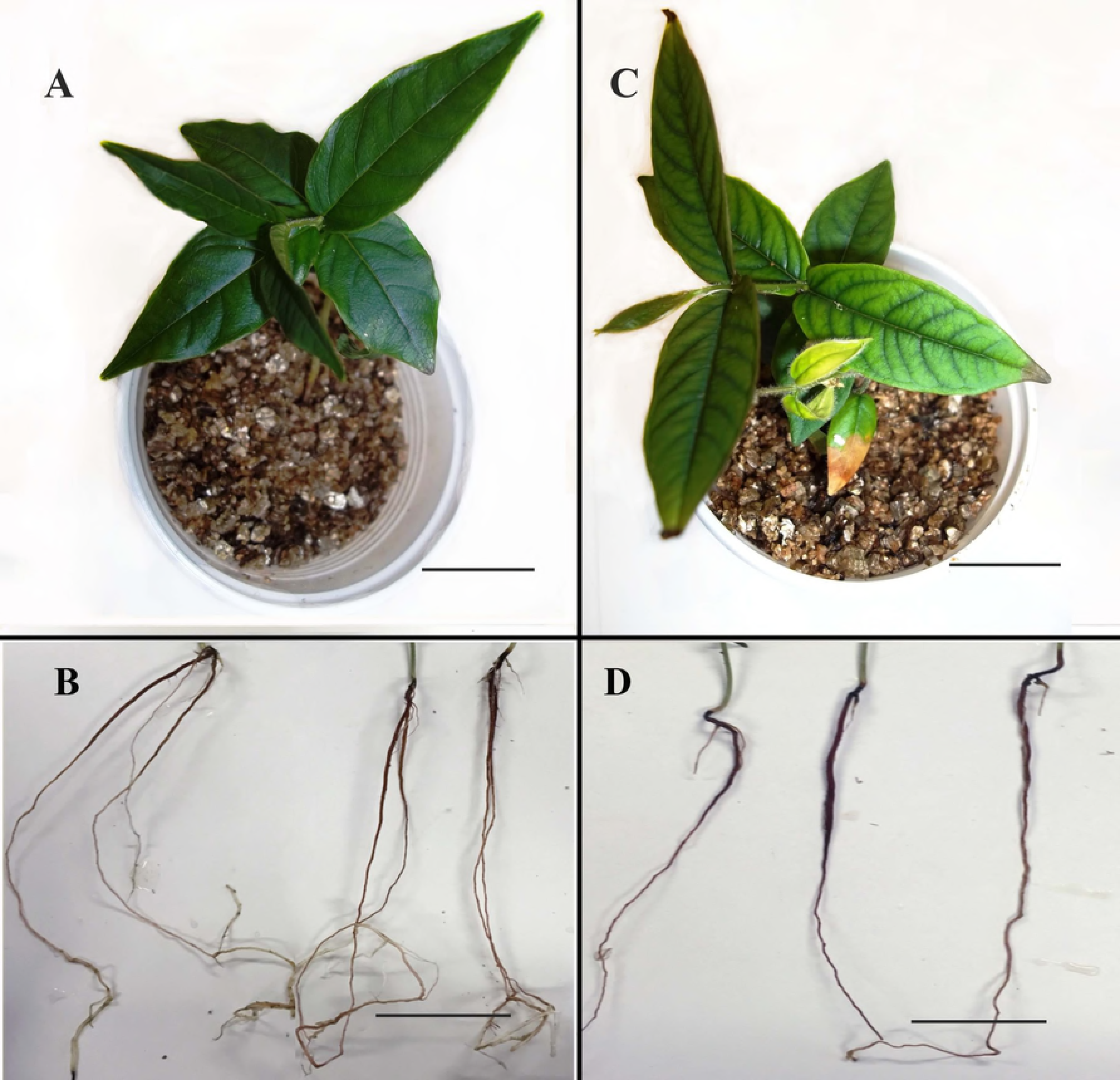
Protein-protein interaction analysis based on STRING with an interaction confidence score (0.900). (A) STRING analysis for differentially abundant proteins upregulated in Al-treated *Q. grandiflora* roots. Red highlighted nodes are ribosomal family proteins. (B) STRING analysis of downregulated differentially abundant proteins in Al-treated *Qualea grandiflora* roots. Red highlighted nodes are proteins related to stress response.

## 4. DISCUSSION

### 4.1. *Qualea grandiflora* required Al for growth and development

Al is harmful to many plant species. Nonetheless, *Q. grandiflora* belongs to a category of plants that requires Al to grow and develop [5, 6]. Although the focus of the current investigation is the proteome of *Q. grandiflora* roots, the data on shoot growth, morphology and ultrastructural chloroplast features are crucial to establish the general health condition of plants from both treatments. The primary observation is that *Q. grandiflora* growth was stimulated by Al. Also, the morphological analysis of shoots from plants that did not receive Al revealed severe signs of stress such as low shoot elongation and chlorosis.

Leaf chlorosis is usually associated with either mineral deficiency [17] or mineral toxicity [18]. In *Q. grandiflora* case, the results showed that the lack of Al negatively affected chlorophylls and carotenes contents, and, at the subcellular level, the chloroplast integrity (Fig. 3). Furthermore, low photosynthetic pigment contents and chloroplast deformities could indicate that an oxidative stress process took place. For instance, the moss *Taxithelium Nepalense*, exposed to toxic amounts of Pb and Cd had lower chlorophyll content and chloroplast abnormalities as well as increased levels of ROS (Reactive Oxygen Species) [19]. Besides, in rice and *Arabidopsis thaliana*, the NADPH thioredoxin reductase (NTRC), a REDOX system, is considered essential to protect chloroplasts [20]. Moreover, it has been shown that *Q. grandiflora* chloroplasts store Al, which does not affect their functionality and structure [6]. Therefore, the current results strongly suggest that Al starvation is highly harmful to *Q. grandiflora* plants.

Concerning roots, the results also support the idea that the absence of Al is a source of stress for *Q. grandiflora*. Based on the morphological and root growth data, this hypothesis appears quite reasonable as the lack of Al made the roots shorter, less branched, and with lower biomass. Additionally, as it is discussed below, these morphological and growth data are coherent with the proteomic analysis of roots since Al induced the upregulation of several metabolic routes, as well as cellular processes and molecular functions that are crucial for plant growth and development.

### 4.2. Proteomic analysis showed that Al treatment upregulated *Q. grandiflora* primary metabolism associated processes favouring root growth and development

Proteomic analysis has been a valuable tool to study the effects of Al exposure in many plants. In *Q. grandiflora*, 410 Al-responsive differentially expressed proteins were identified, which were associated with various biological processes (BP), cellular components (CC) and molecular functions (MF). The majority (62%) of the Al-responsive proteins in *Q. grandiflora* roots were associated with cellular and metabolic processes such as nitrogen compound metabolism, biosynthetic process, catabolism, and primary metabolism. All these processes are crucial during plant growth and development.

Besides, proteins related to cellular components such as ribosomes, cell membranes, mitochondria, and plastids had their abundance increased in Al-treated plants (Fig 5). Conversely, STRING (protein-protein interaction) analysis revealed that stress-related proteins were downregulated in Al-treated *Q. grandiflora* roots (Fig 8). The fact that stress-related pathways were downregulated in plants treated with Al shows that Al did not induce stress in *Q. grandiflora* as it does in many species. Moreover, this fact strengthens the hypothesis that this plant may have an Al-dependent metabolism.

To unveil which metabolic pathways are modulated in *Q. grandiflora* roots by Al, a KEGG analysis was performed to identify proteins and respective metabolic routes responsive to this metal. Thus, Al-responsive metabolic pathways were, to a great extent, associated with primary metabolism components. Consequently, aspects of the primary metabolism that involve carbohydrate metabolism, energy metabolism, lipid metabolism, and aspects of genetic information processing will be further discussed considering their importance of Al for the growth and development of *Q. grandiflora*.

#### 4.2.1. Al induced upregulation of proteins involved in synthesis of cell wall components as well as in the regulation of its properties

Al can interact with various extra and intracellular structures and several mechanisms have been proposed to explain the effects of Al on many plants [21]. Furthermore, the cell wall is the first site to perceive Al presence, which triggers the response to it [22]. Plant Al response will be dependent upon specific characteristics such as sensitivity, tolerance and/or resistance to this metal.

*Q. grandiflora* root proteomic analysis revealed 23 cell wall-related proteins which had their abundance increased in roots of Al-treated plants (Supplementary Table 3). It is remarkable that none of the downregulated proteins was associated with the cell wall in *Q. grandiflora* roots. Consequently, the upregulation of this number of proteins in *Q. grandiflora* roots treated with Al is strongly compatible with the results of root growth analysis presented here. Growing tissues and organs have a high demand for the synthesis and well-structured cell walls.

A comparison between root proteomes from two rice varieties with different levels of Al-tolerance revealed how Al affected cell wall differentiation, and ultimately root growth [22]. When Al-sensitive rice plants (var. Kasalath) were exposed to Al, there was a significant inhibition of root growth [22]. Inversely, the Al-treated tolerant variety (var. Koshihikari) had significantly lower root growth inhibition compared with the sensitive plant [22]. These distinct responses to Al were directly related to the modifications in the abundance of cell wall-related proteins, where some of the proteins were downregulated [22]. Moreover, it was suggested that the differences in abundance of cell wall proteins might have affected the physicochemical properties of the wall, and consequently cell elongation, which was considered a significant Al-related stress response [22]. Hence, the maintenance of cell wall integrity and properties could be considered an essential feature for plant Al tolerance mechanisms, since cell growth and expansion is directly associated with the physicochemical characteristics of the wall [23].

Among the 23 Al responsive cell wall proteins found in *Q. grandiflora* several enzymes play critical roles in cell wall structural integrity, properties, and synthesis. Among the identified proteins there were germin-like proteins, pectinesterases, a fasciclin-like arabinogalactan (8), polygalacturonases, and two alpha-galactosidases (Supplementary Table 3).

For instance, germins and germin-like proteins (GLPs) are ubiquitously found in plants and have been associated with various developmental and biological processes including cell wall deposition and structure [24, 25, 26]. Furthermore, two extra-cellular-matrix associated GLPs were investigated in *Arabidopsis* plants [27]. The authors hypothesized that GLPs proteins might be a class of receptors in the extra-cellular-matrix due to their number of sequences and expression patterns. Therefore, they would be able to respond accordingly to a vast diversity of external environmental factors [27]. If this is true, GLPs could play a role in recognition of Al in the extra-cellular-matrix in *Q. grandiflora* triggering a chain of developmental events in its roots.

Also, pectinesterases are crucial for cell wall integrity. These enzymes control the degree of methyl-esterification of pectins, which is directly associated with cell adhesion and cell wall stiffness [28, 29]. Plant pectins are synthesized highly methyl-esterified, and pectin de-esterification enhances the adhesion properties of the cell wall [28, 29]. Therefore, pectinesterases act in a very controlled manner and regulate the degree of pectin methyl-esterification just after its deposition in the cell wall [29]. This process is critical for plant development, and any disturbance in this process may lead to abnormalities in the cell wall [28].

It has been demonstrated that, in Al-sensitive plants, the accumulation of Al alone may result in increased cell wall stiffness through the cross-linking of pectin methyl residues, which inhibits cell wall loosening required for root growth [30]. Furthermore, in sensitive plants, root tips are the primary sites of Al^3+^ toxicity, and most of the Al in roots are found in the cell wall [30]. This is not observed in *Q. grandiflora*. On the contrary, in this plant, the whole process of root growth and development benefits from Al presence.

The other proteins mentioned above strongly support this statement. Arabinogalactan proteins are present in cell walls and directly involved in plant growth and development, cell adhesion, cell division, and many other developmental processes [31, 32]. For instance, fasciclin-like arabinogalactan proteins were highly present in the cortex and vascular tissues of *Arabidopsis* roots [33]. Fasciclin domains are associated with cell adhesion in many organisms such as plants, fungi, animals, and algae [33, 34]. Besides, lower expression of these proteins in *Arabidopsis* mutants induced the formation of thinner and looser cell walls [33]. It has been speculated that these structural proteins interact with other cell wall components to maintain the structure of the wall [33].

Additionally, two types of proteins found in *Q. grandiflora* roots, the alpha-galactosidases, and polygalacturonases may regulate development by playing a role in cell wall loosening and, thereby, facilitating cell wall expansion [35, 36]. Besides, polygalacturonases play a crucial role in root cap development by degrading demethylated pectin allowing the natural sloughing off outer root cap cells [37]. Root caps are essential for root growth because they protect the root apical meristem as well as facilitates soil penetration and exploration.

Moreover, a putative alpha-galactosidase located at the cell wall of barley leaves was essential for leaf elongation and expansion [35]. In *Arabidopsis* roots, an alpha-galactosidase gene was co-expressed with *AGAMOUS-LIKE42* (*AGL42*), a transcription factor whose expression is proper of root stem cells [38]. This fact might suggest that these enzymes could help to maintain the integrity of root apical meristem, which would favour root growth and development.

It is also worthy to point out that cell wall synthesis is directly dependent on the primary metabolism. Therefore, it is not a coincidence that, in *Q. grandiflora* roots, Al promoted the upregulation of primary metabolic pathways, which is shown by the fact that about 62% of all primary metabolism-related proteins had their abundance increased. Furthermore, KEGG analysis identified 66 proteins with increased relative abundance were involved in carbohydrate metabolism, essential to produce the buildings blocks of cell wall components. Therefore, the increase in cell wall protein abundances in *Q. grandiflora* plants grown with Al indicates an active role of Al in cell wall synthesis and expansion of *Q. grandiflora* roots.

#### 4.2.2. Lipid metabolism in roots of Q. grandiflora

The comparative analysis of differentially abundant proteins in response to Al in *Q. grandiflora* roots revealed 20 proteins involved in lipid metabolism whose abundances increased in response to Al. A KEGG analysis showed that fatty acid degradation pathways were significantly enriched (Table 3). Thus, the most abundant regulated proteins associated to fatty acid degradation were: acyl-CoA oxidase (ACOX), alcohol dehydrogenase (ADH5), acetyl-CoA C-acetyltransferase (atoB), acetyl-CoA C-acyltransferase 1 (ACAA1), enoyl-CoA hydratase/3-hydroxyacyl-CoA dehydrogenase (MFP2) and alcohol class-P dehydrogenase (ADH1).

Consistent with *Q. grandiflora* data, in hydrangea (*H. macrophylla*), also an Al-accumulating plant, a KEGG analysis showed that “lipid metabolism” was significantly enriched in roots and leaves in response to this metal [39]. Thus, the presence of Al increases the abundance of proteins associated with lipid metabolism in hydrangea, which may reflect on plasma membrane functions [39]. Besides, it is believed that the activation of this metabolic pathway in these plants is to maintain plant metabolism and growth by producing energy.

Furthermore, it is noteworthy that enzymes capable of producing substrates for TCA and electron transport chain were upregulated in Al-treated *Q. grandiflora* plants. For instance, the acyl-CoA oxidase (ACOX), an enzyme that catalyses the first step in fatty acid beta-oxidation in peroxisomes and affects various physiological process such as seed oil storage, lipid membrane turnover, senescence, synthesis of some plant hormones, and, consequently plant growth and development [40]. Fatty acid beta-oxidation results in the formation of acetyl-CoA that can be used in the tricarboxylic acid (TCA) cycle. Similarly, the alcohol dehydrogenase (ADH5) was another protein whose abundance increased with Al. Besides, this enzyme belongs to a large family of enzymes that oxidate alcohols to aldehydes and concomitantly reduce NAD^+^ to NADH [41]. Also, it has been established that ADH5 is found in the cytoplasm [41], where it prevents the accumulation of formaldehyde.

Finally, the last example of an enzyme associated with lipid metabolism that might have an impact on *Q. grandiflora* growth and development is the acetyl-CoA C-acetyltransferase (atoB). This enzyme condensates two acetyl-CoA molecules to form acetoacetyl-CoA, which is the first enzymatic step in the biosynthesis of isoprenoids, which comprise a wide range of molecules of high biological significance such as carotenoids, chlorophylls, and plastoquinone. Moreover, isoprenoids are components of the cytokinin phytohormonal class, including, the most common cytokinin, the zeatin [42]. Cytokinins are synthesized in various plant organs; however, one of the main sites of cytokinin synthesis is roots [43]. In fact, until recently, cytokinins were believed to be synthesized exclusively in roots. These hormones affect several physiological and morphogenetic events including cell division, vascular differentiation, apical dominance, senescence, sink and source balance, and consequently nutrient uptake [42]. Therefore, the upregulation of *ACOX, ADH5*, and *atoB* may have contributed to the fact that Al-treated *Q. grandiflora* roots had better growth rate and accumulated more biomass.

### 4.3. Genetic information processing and respective associated cellular components were positively responsive to Al in *Q. grandiflora* roots

It is noteworthy that the KEGG analysis indicated that about 30% of the differentially abundant proteins were related to genetic information processing, which is the highest among all categories catalogued (Fig 7). That is not a coincidence. Concomitantly, elevated levels of ribosome-associated proteins as wells as those related to nucleotide and amino acid metabolisms strengthen the idea of highly intense processing of genetic information. Moreover, enrichment analysis (STRING) of metabolic pathways was coincident with KEGG findings, indicating that ribosomal proteins, amino acid metabolism, and biosynthesis, pyrimidine, and purine metabolism were responsive to Al in roots of *Q. grandiflora* (Table 3).

#### 4.3.1. Nucleotide and amino acid metabolism in roots of Q. grandiflora and Al

Proteins involved in nucleotide metabolism showed altered abundances in response to Al in roots of *Q. grandiflora*. Therefore, four proteins associated with purine anabolic pathways had their abundances significantly increased, and two proteins decreased in response to this metal. Regarding pyrimidine anabolism, five proteins were found with increased abundance in Al-treated plants. This fact may indicate that Al upregulated the synthesis of these nitrogenous bases.

Al is known to have a high affinity for oxygen donor compounds such as inorganic phosphate, nucleotides, RNA, DNA, and proteins. In sensitive plants, the binding of Al to these compounds will result in structural damages, especially in roots, since this organ is the first to have contact with this metal. However, this does not appear to be true for *Q. grandiflora*. The results seem to point to an increase in nucleotide metabolism to likely meet the demands of nucleic acid synthesis and genetic information processing during plant growth.

Plant growth is based on two factors, cell division and cell elongation, which is the principal component of plant growth. Cell elongation requires large amounts of cell wall synthesis, as discussed above. Although cell division also requires cell wall synthesis, to a considerably lesser extent, DNA replication and RNA synthesis are a sine qua non condition for cell division. Furthermore, pyrimidine and purine are the backbones of these nucleic acids. As roots from Al-treated *Q. grandiflora* plants grew and developed significantly better than those from non-treated plants, they need to synthesize these compounds more intensely. This fact may be the basis for pyrimidine and purine related proteins to be positively responsive to Al in *Q. grandiflora* roots.

Concerning amino acids metabolism, the significantly higher abundance of S-adenosylmethionine synthetase (SAMS) may point to an increased synthesis of amino acids [44] and control of gene expression [45], indicating active cell proliferation [46]. Cysteine and aspartate are precursors of methionine synthesis. Besides SAMS, the enzymes malate dehydrogenase (MDH1), MDH2, aspartate semialdehyde dehydrogenase (ASD), and AGXT2 (alanine-glyoxylate transaminase) also had their abundance increased in *Q. grandiflora* in response to Al. The enzyme ASD plays a key role in amino acid synthesis in several organisms, including higher plants [47, 48]. Malate dehydrogenases are enzymes of wide occurrence in living organisms whose principal substrate is the oxaloacetate and are involved in several metabolic pathways, which includes amino acid synthesis [49].

Nonetheless, SAMS might be implicated in metabolic pathways that go beyond amino acid metabolism. This enzyme is necessary for S-adenosylmethionine (SAM) biosynthesis [50]. SAM is the principal methyl donor for the vast majority of transmethylation reactions [51]. These reactions are accomplished by SAM-dependent methyltransferases and crucial for the synthesis of many cellular components such as cell wall, lignin, as well as for RNA capping and epigenetic regulation of gene expression [51, 28]. Thereby, SAMS is directly involved in genetic information processing.

Besides, to the best of our knowledge, the upregulation of SAMS observed in *Q. grandiflora* in response to Al has no parallel in crop plants. For instance, in rice plants, this enzyme was downregulated under Al stress [22]. In addition, in rye and rice, the transcripts and/or protein levels of SAMS also decreased in response to Al [50, 22]. Therefore, the upregulation of SAMS might indicate its involvement in many cellular and metabolic processes that have been induced to cope with *Q. grandiflora* growth and developmental needs. To accomplish that, all processes involved in genetic information processing play essential roles. Consequently, *Q. grandiflora* may require Al to maintain its metabolism functional.

### 4.4. Other metabolic processes responsive to Al in *Q. grandiflora* roots and respective putative role

GO term analysis of *Q. grandiflora* roots showed that several other cellular components were responsive to Al treatment. Among which there were plasma membrane and integral membrane components as well as proteins associated with mitochondria and plastids (Fig 5). Furthermore, the most active molecular functions of proteins were related to catalytic and binding activities (Fig 6). Therefore, a few aspects that involve cellular components and protein molecular functions that have not yet been addressed and are crucial for plant growth and development will be further discussed.

#### 4.4.1. Redox associated enzymes were upregulated in roots of Q. grandiflora plants grown with Al

In *Q. grandiflora*, several redox associated proteins were identified in plants supplemented with Al. Compared with no Al treatment, it was found that 58 and 27 redox associated proteins were up and downregulated, respectively (Supplementary Table 4). For instance, formate dehydrogenase (FDH) was upregulated in *Q. grandiflora* roots. FDHs are versatile enzymes and often associated with response to environmental changes [52]. In *Arabidopsis,* this enzyme is involved in responses to biotic and abiotic stimuli, and its abundance increased in the presence of Al [53]. Moreover, in *Vigna umbellata* this enzyme confers greater tolerance to Al^+3^ [53]. In *V. umbellata*, there was a rapid FDH accumulation in root apices after Al-exposure. Furthermore, in this plant Al^+3^ was considered the inducer of *VuFDH* expression [53].

FDHs catalyse the oxidation of formate to carbon dioxide resulting in the formation of NADH. Moreover, in plants, these enzymes are localized in mitochondria and supply organisms with energy and reducing power in the form of ATP [54, 55]. Also, formate oxidation by FDHs may be coupled with the oxidative phosphorylation [56]. Compared with no Al-treatment, Al supplementation was an abiotic stimulus for *Q. grandiflora* plants, instead of a stress eliciting agent. Therefore, FDHs might help to keep the energy supply steady, which can explain the higher biomass accumulation of Al-treated plants. Additionally, the upregulation of proteins such as FDH may also have conferred an evolutive advantage to these plants, which could have contributed to the adapt *Q. grandiflora* to the Cerrado edaphic conditions.

Another class of proteins that in *Q. grandiflora* had their abundance significantly altered in response to Al was the antioxidant enzymes that play a role against oxidative stress. In *Q. grandiflora,* most antioxidant enzymes had their abundance diminished. Conversely, in roots of sensitive tomato plants exposed to Al, proteins such as monodehydroascorbate reductase (MDHAR), superoxide dismutase (SOD), and selenium-proteins (not found in *Q. grandiflora*), all with redox activities were positively regulated [57]. Also, the disulphide isomerase (PDI), a protein that repairs denatured proteins was also upregulated in sensitive tomato plants [57]. Likewise, in rice, which is one of the most Al tolerant cultures, the antioxidant proteins SOD, glutathione S-transferase (GST), and S-adenosylmethionine synthetase 2 (SAMS) are consistently induced by Al [50]. Thereby, Al tolerance is a complex phenomenon that involves multiple proteins, distinct biological processes and a broad variety of metabolic response. As discussed, the metabolic response may differ from species to species. Nevertheless, either in sensitive or tolerant plants, it appears that Al induces oxidative stress response mechanisms. In *Q. grandiflora* case, our data support the idea that Al is not a stress elicitor.

#### 4.4.2. Al affected organic acids metabolism in Q. grandiflora

In *Q. grandiflora* roots, it was observed a possible upregulation of glyoxylate metabolism due to an increased abundance of proteins such as malate dehydrogenase 1 and 2 (MDH1/MDH2), formate dehydrogenase (FDH), catalase (CAT), acetyl-CoA C-acetyltransferase (atoB), serine hydroxymethyltransferase (SHMT) and glycine H cleavage system (GCSH) in plants treated with Al. These results suggest that oxalate biosynthesis in roots of Al grown *Q. grandiflora* plants was active and may assist in the capture and accumulation of this metal in this plant.

Organic acids such as oxalate and citrate have often been involved in the process of tolerance and accumulation of Al in various plants, such as wheat and buckwheat [58, 59, 60]. In plants, there are three routes for oxalate biosynthesis: the glyoxylate, the ascorbate, and the oxaloacetate pathways [61, 62]. For example, in rice, glyoxylate was an efficient precursor for oxalate biosynthesis, which may be a product of the glyoxylate cycle [62].

It is important to note that there are studies that correlate organic acids to chelation and Al-transportation. Hence, oxalate can chelate Al^3+^ and form with it a stable complex. Plants release organic acids to prevent Al-toxicity, both externally (rhizosphere) and internally [63, 59, 60]. In buckwheat, an Al-tolerant species, when oxalate secretion is suppressed by an anion channel inhibitor, e.g., phenylglyoxal, root elongation is inhibited in the presence of Al [64, 65]. This fact demonstrates the role of organic acids in response to Al in plants. This phenomenon is also seen in other plants such as Hydrangea macrophylla [39].

Additionally, in *Melastoma malabathricum* L. (Melastomataceae) Al transportation is suggested to take place in the xylem, probably in the form of Al-citrate complexes [66]. The type of organic acid involved in Al response may vary with the species. For instance, *Triticum aestivum* L. secretes malate [58], *Phaseolus vulgaris* L. [67], *Zea mays* L. [68] and *Fagopyrum esculentum* Moench exudates oxalate in response to Al [69].

As mentioned, in *Q. grandiflora* the increases in abundance of several enzymes of this glyoxylate route have been observed. It seems that the increase of oxalate biosynthesis in Al-accumulating plants is a default component in response to Al. Although there are no studies on the function of these organic acids in *Q. grandiflora* roots, these compounds might be involved in the capture and transportation of Al in this species. Nonetheless, a metabolic profile of *Q. grandiflora* may help to determine the function of these compounds in this plant.

Another possibility involves the upregulation of malate dehydrogenases (MDH) in plants treated with Al. There are several MDH isoforms in plants, which have been located in various cellular compartments such as cytoplasm, peroxisomes, chloroplasts, and mitochondria [70]. Furthermore, these enzymes play many physiological functions. For instance, in mitochondria, mMDH plays a role in TCA cycle where it oxidizes malate to OAA to form citrate [71]. Al stimulates the synthesis and root efflux of citrate in several plant species [72, 73]. It has been hypothesized that Al is transported to shoots as Al-citrate complexes through xylem vessels [74], and, maybe, citrate could play a role in Al transportation in *Q. grandiflora* as well. It is interesting that a metabolomic analysis of *Q. grandiflora* roots has shown a citrate content significantly higher in Al-treated plants (Silva, unpublished results). This matter is currently under investigation.

Besides, MDH isoforms are involved in the transport and utilization of malate and OAA as well as in the availability of NAD in several organelles [71]. Additionally, Wang et al. (2015) associated the high expression of an MDH (GhmMDH1) in transgenic cotton plants with a more efficient uptake of insoluble phosphorus (Al-P, Fe-P, or Ca-P). *GhmMDH1*-overexpressing cotton plants that were supplied exclusively with these forms of insoluble P developed longer roots with higher biomass than wild-types plants [71].

This fact is highly relevant because Al-treated *Q. grandiflora* plants had longer roots and higher root biomass as well. As mentioned previously, Cerrado soils are acid and abundant in Al, and thereby, have high levels of insoluble P. It is possible that these edaphic conditions were a significant factor in the evolutive process of native species. Therefore, there was a selective pressure that favoured plant adaptations to deal with these harsh environmental conditions. Consequently, the *Q. grandiflora* MDHs might have evolved to work with insoluble P due to Cerrado soils properties. It is not inconsistent to think that, maybe, these plants have MDHs that work better in the presence of Al than without it. Nonetheless, this is another exciting hypothesis that requires testing.

## 5. A Model Proposal for Biochemical and Physiological Cellular Processes Upregulated by Al in *Q. grandiflora* Root Cells

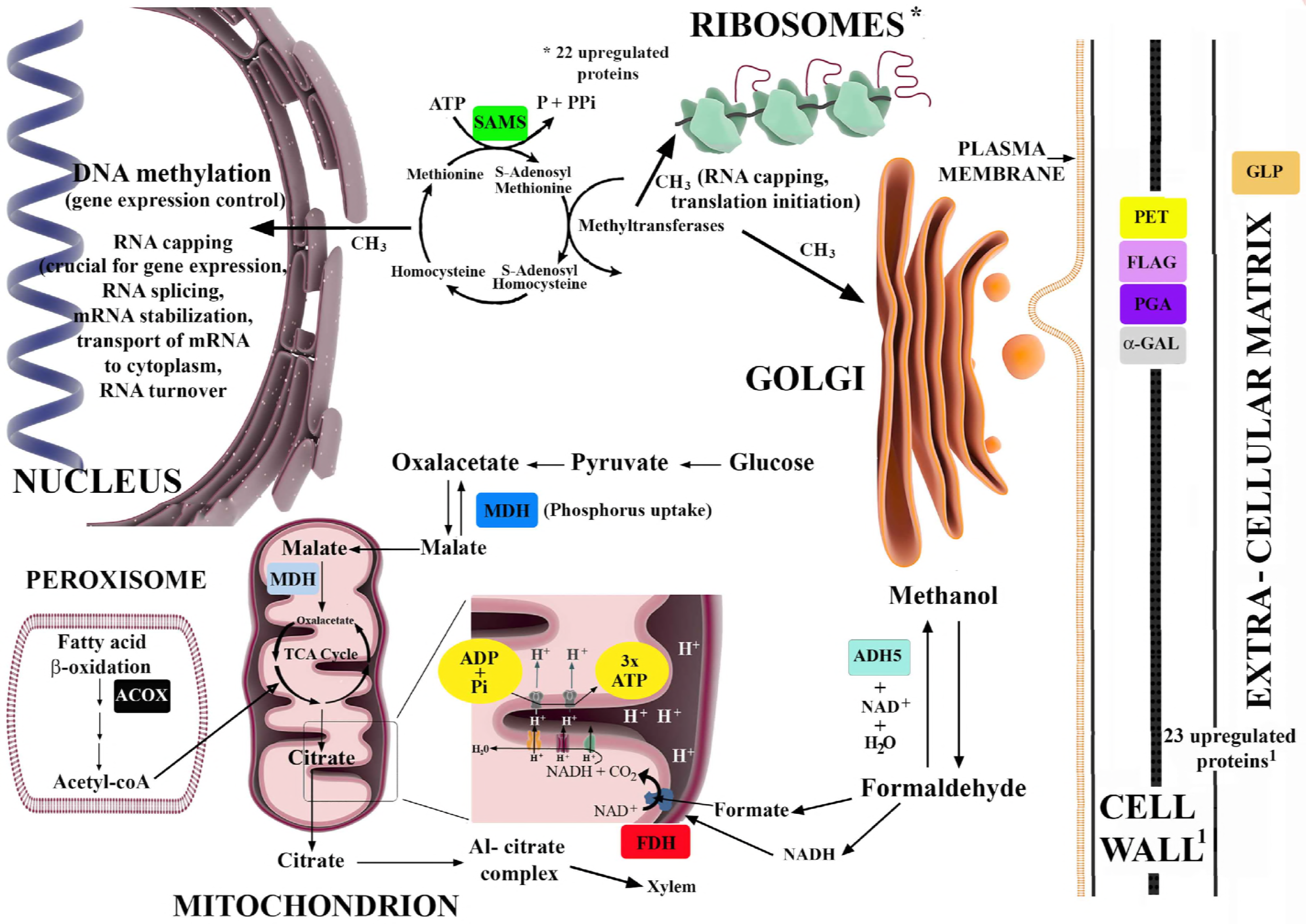

### Extra-Cellular Matrix

Germin-like proteins (GLP): some GLP are believed to be a class of receptors in the extra-cellular-matrix.

### Cell Wall

#### Proteins

1. Pectinesterases (PET); 2. Fasciclin-like arabino galactosidase (FLAG); 3. Polygalacturonases (PGA); 4. Alpha-galactosidase (α-GAL).

#### Physiological and Metabolic Events

These proteins may be involved in several processes such as loosening the cell wall (Pectinesterases), which facilitates the cell expansion and elongation (Pectinesterases, alpha-galactosidase), work as Al receptor (Germin-like proteins), root cap development and integrity (Polygalacturonases), play a role in stem cells of root apical meristem (alpha-galactosidase).

### Cytoplasm

#### Proteins

S-Adenosyl Methionine Synthetase (SAMS) – Essential for the synthesis of S-adenosyl methionine (SAM), which is the main methyl donor crucial for the synthesis of a great number of compounds.

- Carbohydrates components of pectins. These pectin components are synthesized highly methylated, otherwise they would prevent cell expansion due to crosslinking of de-esterified pectin polymers by Ca. Pectin de-esterification is a highly controlled process that involves cell-wall based enzymes named pectinesterases, and it is directly associated with cell wall adhesion/loosening.
- SAMS is believed to be crucial for genetic information processing, as it may provide methyl groups through SAM synthesis for DNA methylation, which is one epigenetic mechanism that often modify gene expression. Besides, SAM is also associated with RNA capping, RNA splicing, RNA transportation, mRNA stability, and translation initiation. Without SAMS, the flow of genetic information would be seriously impaired.

Ribosomes: 22 ribosomal-associated proteins were upregulated, which may positively affect translation of mRNAs, and, consequently protein synthesis.

Malate Dehydrogenases (MDHs) – They could play a role in insoluble phosphorus (P) uptake, which is main form of P in Cerrado Soils. Phosphorus is an essential component of ATP, proteins, and nucleic acids.

Alcohol dehydrogenase (ADH5) – a cytoplasmic enzyme whose activity may result in the production of formate and NADH, regularly used in the electron transport chain to produce ATP.

### Mitochondrion

#### Mitochondrial matrix Proteins

Malate Dehydrogenases (MDHs) – Also, these enzymes could be associated with Al transportation to aerial plant parts through citrate formation. Al-citrate complex might be the form of Al that is translocated throughout the plant by the xylem.

#### Mitochondrial membrane-based protein

FDH – It has a transmembrane domain in the beta subunit, which allows the conduction of electrons within the protein. It catalyses the reaction: Formate + NAD^+^ ⇌ CO_2_ + NADH + H^+^

#### Peroxisomes

##### Protein

Acyl-CoA oxidase (ACOX) – catalyses the first step of beta-oxidation of fatty acids, likely in peroxisomes, which may form acetyl-CoA, an important compound of TCA cycle, and thereby, crucial for ATP production.

## 6. Conclusions

The present work indicates that Al is a required for the growth and development of *Q. grandiflora*. LC-MS-MS analyses provided a comprehensive inventory of Al-responsive proteins in *Q. grandiflora* roots. Principal component analysis (PCA) revealed that Al-treatment induced systemic changes in the proteome of *Q. grandiflora* roots. Bioinformatic analysis revealed a link between changes in protein abundance and various metabolic pathways in response to Al, as well as interactions between these differentially regulated proteins.

*Q. grandiflora* proteome revealed that Al was necessary for the metabolism of this species and directly involved in: a) enhancement of primary metabolism – cell wall synthesis, methylation cycle, protein synthesis; b) increased genetic information processing; c) redox activity; and, d) organic acid metabolism. These results provide valuable insights into the molecular mechanism of native Al-accumulating plants as well as their way of dealing with it. Moreover, the current investigation has expanded knowledge on this topic and enhanced our toolkit for the genetic improvement of cultivated plants more resistant to Al.

Although no conclusion can be drawn about the essentiality of Al in *Q. grandiflora* metabolism, the data supports the idea that this metal has a metabolic role to play in this plant. Therefore, *Q. grandiflora* does resent Al starvation, which leads to a proposal of an Al-dependent metabolism for this species. Nonetheless, the level of dependency has yet to be determined.

## 7. Acknowledgments

We would like to thank the Coordination for the Improvement of Coordination for the Improvement of Higher Level -or Education-Personnel (CAPES), the Brazilian Council of Research (CNPq), and the Support Research of the Federal District Foundation (FAP-DF) for their financial support of this research. Furthermore, we also thank Dr. Fernando Araripe Gonçalves Torres for his valuable help.

## Acknowledgments

We would like to thank the Coordination for the Improvement of Coordination for the Improvement of Higher Level -or Education-Personnel (CAPES) for three graduate student scholarships and a Post-doc scholarship, and the Support Research of the Federal District Foundation (FAP-DF) grant 0193.000941/2015 for their support of this research. The funders had no role in study design, data collection and analysis, decision to publish, or preparation of the manuscript

## Abbreviations

Al: Aluminium
TEM: Transmission Electron Microscopy
MS: Murashige and Skoog medium
LC-MS/MS: liquid chromatography and mass spectrometry

## References

1. Lopes AS, Cox FR. Cerrado vegetation in brazil: an edaphic gradient. Agronomy J. 1977 Sept; 69: 828–831. doi:10.2134/agronj1977.00021962006900050025x.

2. Furley PA, Ratter JA. Soil Resources and Plant Communities of the Central Brazilian Cerrado and Their Development. J. Biogeography. 1988 Jan; 15: 97–108. doi:10.2307/2845050.

3. Haridasan M. Aluminium accumulation by some Cerrado native species of central Brazil. Journal. Plant and soil. 1982; 65: 265–273.

4. Andrade LRM, Barros LMG, Echevarria GF, Amaral LIV, Cotta MG, Rossatto DR, et al. Al-hyperaccumulator Vochysiaceae from the Brazilian Cerrado store aluminum in their chloroplasts without apparent damage. Environ. Exp. Bot. 2011 May 27; 70: 37–42. doi:10.1016/j.envexpbot.2010.05.013.

5. Landim MI, Hingst-Zaher E. Brazil’s biodiversity crisis: natural history collections are vital to preserving Brazil’s biomes. Icomnews. 2010; 2: 14–15.

6. Myers N, Mittermeier RA, Mittermeier CG, Fonseca GA, Kent J. Biodiversity hotspots for conservation priorities. Nature. 2000; 403: 853–858. doi:10.1038/35002501.

7. Delhaize E, Ma JF, Ryan PR. Transcriptional regulation of aluminium tolerance genes, Trends Plant Sci. 2012; 17: 341–348. doi:10.1016/j.tplants.2012.02.008.

8. Iuchi S, Koyama H, Iuchi A, Kobayashi Y, Kitabayashi S, Kobayashi Y, et al. Zinc finger protein STOP1 is critical for proton tolerance in *Arabidopsis* and coregulates a key gene in aluminum tolerance. PNAS. 2007; 104: 9900–9905. doi:10.1073/pnas.0700117104.

9. Sawaki Y., Iuchi S., Kobayashi Y., Kobayashi Y., Ikka T., Sakurai N., et al. STOP1 regulates multiple genes that protect *Arabidopsis* from proton and aluminum toxicities, Plant Physiol. 2009; 150: 281–294. doi:10.1104/pp.108.134700.

10. Hoekenga OA, Maron LG, Piñeros MA, Cançado GMA, Shaff J, Kobayashi Y, et al. *AtALMT1*, which encodes a malate transporter, is identified as one of several genes critical for aluminum tolerance in *Arabidopsis*. PNAS. 2006; 103: 9738–9743. doi:10.1073/pnas.0602868103.

11. Sivaguru M, Ezaki B, He Z.-H, Tong H, Osawa H, Baluska F, et al. Aluminum-induced gene expression and protein localization of a cell wall-associated receptor kinase in *Arabidopsis*. Plant Physiol. 2003; 132: 2256–2266.

12. Mangeon A, Pardal R, Menezes-Salgueiro AD, Duarte GL, Seixas R, Cruz FP, et al. AtGRP3 is implicated in root size and aluminum response pathways in *Arabidopsis*. PLOS One. 2016 Mar 3; 1–11. doi:10.1371/journal.pone.0150583.

13. Wang ZQ, Xu XY, Gong QQ, Xie C, Fan W, Yang JL, et al. Root proteome of rice studied by iTRAQ provides integrated insight into aluminum stress tolerance mechanisms in plants. J Proteom. 2014; 98: 189–205. doi:10.1016/j.jprot.2013.12.023.

14. Murashige T, Skoog F., A revised medium for rapid growth and bioassays with tobacco tissue cultures, Physiol. Plant. 1962; 15: 473–497.

15. Kanehisa M, Sato Y, Morishima K. BlastKOALA and GhostKOALA: KEGG tools for functional characterization of genome and metagenome sequences. J. Mol. Biol. 2016; 428: 726–731. doi:10.1016/j.jmb.2015.11.006.

16. Szklarczyk D, Franceschini A, Wyder S, Forslund K, Heller D, Huerta-Cepas J, et al. STRING v10: protein–protein interaction networks, integrated over the tree of life. Nucleic Acids Res. 2015; 43: D447–D452. doi:10.1093/nar/gku1003.

17. Tewari RK, Kumar P, Neetu, Sharma PN. Signs of oxidative stress in the chlorotic leaves of iron starved plants. Plant Sci. 2005; 169: 1037–1045. doi:10.1016/j.plantsci.2005.06.006.

18. Saglam A, Yetissin F, Demiralay M, Terzi R. Copper stress and responses in plants. In: Plant Metal Interaction, Ed. Parvaiz Ahmad. Elsevier. 2016. pp 21–40. doi. org/10.1016/B978-0-12-803158-2.00002-3.

19. Choudhury S, Panda SK. Toxic effects, oxidative stress and ultrastructural changes in moss *Taxithelium Nepalense* (Schwaegr.) Broth. under chromium and lead phytotoxicity. Water Air Soil Pollut. 2005; 167: 73–90. http://doi.org/10.1007/s11270-005-8682-9.

20. Pérez-Ruiz JM, Spínola MC, Kirchsteiger K, Moreno J, Sahrawy M, Cejudo FJ. Rice NTRC is a high-efficiency redox system for chloroplast protection against oxidative damage. Plant Cell. 2006; 18: 2356–2368. doi/10.1105/tpc.106.041541.

21. Kochian LV, Piñeros MA, Hoekenga OA. The physiology, genetics and molecular biology of plant aluminum resistance and toxicity, Plant and Soil. 2005; 274: 175–195. doi:10.2307/24129042.

22. Wang CY, Shen RF, Wang C, Wang W. Root protein profile changes induced by Al exposure in two rice cultivars differing in Al tolerance. J. Proteom. 2013; 281–293. doi:10.1016/j.jprot.2012.09.03.

23. Cosgrove DJ. Growth of the plant cell wall. Nat. Rev. Mol. Cell Biol. 2005 Nov 1; 6: 850–861. doi:10.1038/nrm1746.G.

24. Zimmermann G, Bäumlein H, Mock H-P, Himmelbach A, Schweizer P. The multigene family encoding germin-like proteins of barley. Regulation and function in basal host resistance. Plant Physiol. 2006; 142: 181–192. doi:10.1104/pp.106.083824.

25. Lu M, Han Y-P, Gao J-G, Wang X-J, Li W-B. Identification and analysis of the germin-like gene family in soybean. BMC Genomics. 2010 Nov 8; 11: 620. doi:10.1186/1471-2164-11-620.

26. Bosch M, Mayer C-D, Cookson A, Donnison IS. Identification of genes involved in cell wall biogenesis in grasses by differential gene expression profiling of elongating and non-elongating maize internodes. J. Exp. Bot. 2011 Mar 14; 62: 3545–3561. doi:10.1093/jxb/err045.

27. Membré N, Bernier F, Staiger D, Berna A. *Arabidopsis thaliana* germin-like proteins: common and specific features point to a variety of functions. Planta. 2000; 211: 345–354. doi:10.1007/s004250000277.

28. Pereira LAR, Todorova M, Cai X, Makaroff CA, Emery RJN, Moffatt BA. Methyl recycling activities are co-ordinately regulated during plant development. J. Exp. Bot. 2007; 58: 1083–1098. doi:10.2307/24036568.

29. Kohli P, Kalia M, Gupta R. Pectin methylesterases: a review. J Bioprocess Biotech. 2015; 5:227 1000227 doi:10.4172/2155-9821.100022.

30. Tabuchi A, Matsumoto H. Changes in cell-wall properties of wheat (*Triticum aestivum*) roots during aluminum-induced growth inhibition. Physiol. Plant. 2001; 112: 353–358. doi:10.1034/j.1399-3054.2001.1120308.x.

31. Johnson KL, Jones BJ, Bacic A, Schultz CJ. The fasciclin-like arabinogalactan proteins of *Arabidopsis*. A multigene family of putative cell adhesion molecules. Plant Physiol. 2003; 133: 1911–1925. doi:10.1104/pp.103.031237.

32. Zang L, Zheng T, Chu Y, Ding C, Zhang W, Huang Q, Su X. Genome-wide analysis of the fasciclin-like arabinogalactan protein gene family reveals differential expression patterns, localization, and salt stress response in *Populus*. Front Plant Sci. 2015; 6: doi:10.3389/fpls.2015.01140.

33. Shi H, Kim Y, Guo Y, Stevenson B, Zhu J-K. The *Arabidopsis SOS5* locus encodes a putative cell surface adhesion protein and is required for normal cell expansion. Plant Cell. 2003; 15: 19–32. doi.org/10.1105/tpc.007872.

34. Ames RM. Using network extracted ontologies to identify novel genes with roles in appressorium development in the rice blast fungus *Magnaporthe oryzae*. Microorganisms. 2017 Jan 17; 5. doi:10.3390/microorganisms5010003.

35. Chrost B, Kolukisaoglu U, Schulz B, Krupinska K. An α-galactosidase with an essential function during leaf development. Planta. 2007 Jul 15; 225: 311–320. doi:10.1007/s00425-006-0350-9.

36. Minic Z. Physiological roles of plant glycoside hydrolases. Planta. 2008; 227: 723–740. doi:10.1007/s00425-007-0668-y.

37. Kamiya M, Higashio S-Y, Isomoto A, Kim J-M, Seki M, Miyashima S, Nakajima K. Control of root cap maturation and cell detachment by BEARSKIN transcription factors in *Arabidopsis*. Development. 2016 Nov 1; 143: 4063–4072. doi:10.1242/dev.142331.

38. Rodriguez RE, Ercoli MF, Debernardi JM, Breakfield NW, Mecchia MA, Sabatini M, et al. MicroRNA miR396 regulates the switch between stem cells and transit-amplifying cells in *Arabidopsis* roots. Plant Cell. 2015; 27: 3354–3366. doi:10.1105/tpc.15.00452.

39. Chen H, Lu C, Jiang H, Peng J. Global transcriptome analysis reveals distinct aluminum-tolerance pathways in the Al-accumulating species *Hydrangea macrophylla* and marker identification. PLoS One. 2015 Dec 14; 10 e0144927. doi:10.1371/journal.pone.0144927.

40. Arent S, Pye VE, Henriksen A. Structure and function of plant acyl-CoA oxidases, Plant Physiol. Biochem. 2008 Jan 3: 46: 292–301. doi:10.1016/j.plaphy.2007.12.014.

41. Smidt O, du Preez JC, Albertyn J. The alcohol dehydrogenases of *Saccharomyces cerevisiae*: a comprehensive review. FEMS Yeast Res. 2008; 8: 967–978. doi:10.1111/j.1567-1364.2008.00387.x.

42. Kieber JJ, Schaller GE. Cytokinins. In The Arabidopsis Book. 2. 2014. 12:e0168. doi:10.1199/tab.0168.

43. Xu Y, Burgess P, Zhang X, Huang B. Enhancing cytokinin synthesis by overexpressing ipt alleviated drought inhibition of root growth through activating ROS-scavenging systems in *Agrostis stolonifera*. J. Exp. Bot. 2016; 67: 1979–1992. doi:10.1093/jxb/erw019.

44. Markham GD, Pajares MA. Structure-function relationships in methionine adenosyltransferases. Cell. Mol. Life Sci. 2009 Oct 27; 66: 636–648. doi:10.1007/s00018-008-8516-1.

45. Reytor E, Pérez-Miguelsanz J, Alvarez L, Pérez-Sala D, Pajares MA. Conformational signals in the C-terminal domain of methionine adenosyltransferase I/III determine its nucleocytoplasmic distribution. FASEB J. 2009; 23: 3347–3360. doi:10.1096/fj.09-130187.

46. Yoon S, Lee W, Kim M, Kim TD, Ryu Y. Structural and functional characterization of S-adenosylmethionine (SAM) synthetase from *Pichia ciferrii*. Bioprocess Biosyst Eng. 2012; 35: 173–181. doi:10.1007/s00449-011-0640-x.

47. Jander G, Joshi V. Aspartate-derived amino acid biosynthesis in *Arabidopsis thaliana*. In Arabidopsis Book. 2009. 7e0121. doi:10.1199/tab.0121.

48. Viola RE, Faehnle CR, Blanco J, Moore RA, Liu X, Arachea BT, Pavlovsky AG. The catalytic machinery of a key enzyme in amino acid biosynthesis. J. Amino Ac. 2011. doi:10.4061/2011/352538.

49. Musrati RA, Kollárová M, Mernik N, Mikulásová D. Malate dehydrogenase: distribution, function and properties, Gen. Physiol. Biophys. 1998; 17: 193–210.

50. Yang Q, Wang Y, Zhang J, Shi W, Qian C, Peng X. Identification of aluminum-responsive proteins in rice roots by a proteomic approach: cysteine synthase as a key player in Al response. Proteomics. 2007; 7: 737–749. doi:10.1002/pmic.200600703.

51. Moffatt BA, Weretilnyk EA. Sustaining S-adenosyl-l-methionine-dependent methyltransferase activity in plant cells. Physiol. Plant. 2001; 113: 435–442. doi:10.1034/j.1399-3054.2001.1130401.x.

52. Jormakka M, Byrne B, Iwata S. Formate dehydrogenase–a versatile enzyme in changing environments. Curr. Opin. Struct. Biol. 2003; 13: 418–423.

53. Smidt O, du Preez JC, Albertyn J. The alcohol dehydrogenases of *Saccharomyces cerevisiae*: a comprehensive review. FEMS Yeast Res. 2008; 8: 967–978. doi:10.1111/j.1567-1364.2008.00387.x.

54. Popov VO, Lamzin VS. NAD(+)-dependent formate dehydrogenase. Biochem J. 1994; 301: 625–643.

55. Alekseeva AA, Savin SS, Tishkov VI. NAD +-dependent formate dehydrogenase from plants. Acta Naturae. 2011 Oct-Dec; 3: 38–54.

56. Oliver DJ. Formate oxidation and oxygen reduction by leaf mitochondria. Plant Physiol. 1981; 68: 703–705.

57. Zhou S, Okekeogbu I, Sangireddy S, Ye Z, Li H, Bhatti S, et al. Proteome modification in tomato plants upon long-term aluminum treatment. J. Proteome Res. 2016; 15: 1670–1684. doi:10.1021/acs.jproteome.6b00128.

58. Delhaize E, Craig S, Beaton CD, Bennet RJ, Jagadish VC, Randall PJ. Aluminum tolerance in wheat (*Triticum aestivum* L.) (I. uptake and distribution of aluminum in root apices). Plant Physiol. 1993; 103: 685–693.

59. Ma JF, Ryan PR, Delhaize E. Aluminium tolerance in plants and the complexing role of organic acids. Trends Plant Sci. 2001 Jun 1; 6: 273–278.

60. Yokosho K, Yamaji N, Ma JF. Global transcriptome analysis of Al-induced genes in an Al-accumulating species, common buckwheat (*Fagopyrum esculentum* Moench). Plant Cell Physiol. 2014; 55: 2077–2091. doi:10.1093/pcp/pcu135.

61. Franceschi VR, Nakata PA. Calcium oxalate in plants: formation and function. Annu. Rev. Plant Biol. 2005 Mar 31; 56: 41–71. doi:10.1146/annurev.arplant.56.032604.144106.

62. Yu L., Jiang J., Zhang C., Jiang L., Ye N., Lu Y., et al. Glyoxylate rather than ascorbate is an efficient precursor for oxalate biosynthesis in rice. J. Exp. Bot. 2010; 61: 1625–1634. doi:10.1093/jxb/erq028.

63. Ma J.F. Role of organic acids in detoxification of aluminum in higher plants. Plant Cell Physiol. 2000; 41: 383–390.

64. Zheng SJ, Ma JF, Matsumoto H. High aluminum resistance in buckwheat. Plant Physiol. 1998; 117: 745–751.

65. Klug B, Horst WJ. Spatial characteristics of aluminum uptake and translocation in roots of buckwheat (*Fagopyrum esculentum*). Physiol. Plant. 2010 Jun 1; 139: 181–191. doi:10.1111/j.1399-3054.2010.01355.x.

66. Watanabe T, Osaki M. Mechanisms of adaptation to high aluminum condition in native plant species growing in acid soils: a review. Comm. Soil Sci. Plant An. 2002; 33: 1247–1260. doi:10.1081/CSS-120003885.

67. Miyasaka SC, Buta JG, Howell RK, Foy CD. Mechanism of aluminum tolerance in snapbeans. Plant Physiol. 1991; 96: 737–743.

68. Pellet DM, Grunes DL, Kochian LV. Organic acid exudation as an aluminum-tolerance mechanism in maize (*Zea mays* L.), Planta. 1995; 196: 788–795. doi:10.1007/BF01106775.

69. Ma JF, Hiradate S, Nomoto K, Iwashita T, Matsumoto H. Internal detoxification mechanism of Al in *Hydrangea*: identification of Al form in the leaves. Plant Physiol. 1997; 113: 1033–1039. doi:10.2307/4277625.

70. Gietl C. Malate dehydrogenase isoenzymes: cellular locations and role in the flow of metabolites between the cytoplasm and cell organelles. Biochim. Biophys. Acta. 1992; 1100: 217–234.

71. Wang Z-A, Li Q, Ge X-Y, Yang C-L, Luo X-L., Zhang A-H, et al. The mitochondrial malate dehydrogenase 1 gene *GhmMDH1* is involved in plant and root growth under phosphorus deficiency conditions in cotton. Sci. Rep. 2015; 5: 10343. doi:10.1038/srep10343.

72. de la Fuente JM, Ramírez-Rodríguez V, Cabrera-Ponce JL, Herrera-Estrella L. Aluminum tolerance in transgenic plants by alteration of citrate synthesis. Science. 1997 Jun 6; 276: 1566–1568.

73. Yang JL, Zhang L, Li YY, You JF, Wu P, Zheng SJ, Citrate Transporters play a critical role in aluminium-stimulated citrate efflux in rice bean (*Vigna umbellata*) roots. Ann Bot. 2006; 97: 579–584. doi:10.1093/aob/mcl005.

74. Mariano ED, Jorge RA, Keltjens WG, Menossi M. Metabolism and root exudation of organic acid anions under aluminium stress. Braz. J. Plant Physiol. 2005; 17: 157–172. doi:10.1590/S1677-04202005000100013.

